# Precision targeting of autoreactive B cells in systemic lupus erythematosus using anti-9G4 idiotope synthetic immune receptor T cells

**DOI:** 10.1101/2025.10.19.682634

**Authors:** Jin Liu, Yuanxuan Xia, Brian J. Mog, Colin Gliech, Elana Shaw, Dylan Ferris, Abigail McGahan, Kennedy A Cleage, Brock Moritz, Kyle J. Kaeo, Tolulope O. Awosika, Sarah R. DiNapoli, Stephanie Glavaris, Xuyang Li, Nikita Marcou, Alexander H. Pearlman, Taha S. Ahmedna, Regina Bugrovsky, Scott A. Jenks, Iñaki Sanz, Chetan Bettegowda, Suman Paul, Victoria Duarte Alvarado, Denis Wirtz, Daniel Goldman, Michelle Petri, Kenneth W. Kinzler, Shibin Zhou, Felipe Andrade, Bert Vogelstein, Maximilian F. Konig

## Abstract

Chimeric antigen receptor (CAR)-T cell therapies can induce drug-free remission in systemic lupus erythematosus (SLE), but indiscriminate B cell targeting causes immunosuppression, unnecessary infections, and cytokine toxicities that preclude widespread use. Here, we overcome this by targeting the 9G4 idiotope, a shared structural feature of pathogenic B cell receptors encoded by the *IGHV4-34* gene. We engineered anti-9G4 CAR-T cells and chimeric TCR-T cells to selectively eliminate autoreactive B cells while preserving protective immunity. Both platforms eradicated autoreactive B cells and autoantibodies in vitro and in vivo, spared normal B cells, and markedly reduced cytokine release compared to conventional CAR-T cells. This precision extended to cold agglutinin disease and lymphoma. These findings establish a framework for IGHV idiotope-directed cellular therapies for treating autoimmune and neoplastic diseases while preserving immune competence.

## Introduction

Systemic lupus erythematosus (SLE) is a potentially life-threatening multisystem autoimmune disease that predominantly affects women of childbearing age (*1*). Despite important therapeutic advances, achieving disease remission in SLE remains challenging (*2*), and uncontrolled disease activity often results in irreversible end-organ damage (*1*). Currently available drugs are broadly immunosuppressive and have narrow therapeutic indices, leading to infections – a major cause of mortality in SLE – and other dose-limiting toxicities (*3*). Therapies that control disease without impairing protective immune responses are critically needed, yet such therapies remain elusive.

Antinuclear antibodies are a canonical feature of SLE (*4*), mediate tissue damage (*5*), and can precede the onset of clinical symptoms by years (*6*). As sources of pathogenic autoantibodies, targeting B cells and plasma cells has been a major therapeutic focus (*7*). The use of monoclonal antibodies targeting CD20+ B cells has shown limited efficacy in SLE (*8–11*), achieving only incomplete depletion of B cells in lymphoid organs and autoimmune target tissues (*12, 13*). By contrast, recent successes in repurposing more potent therapeutic modalities targeting B cells, including bispecific T cell engagers and chimeric antigen receptor (CAR)-T cells, have confirmed the transformative potential of B cell-directed therapies for the treatment of lupus (*14–16*). Autologous CD19 CAR-T-cells, which can effectively deplete B cells in relevant tissue niches (*13*), can achieve normalization of most autoantibody levels and complete – and in some cases durable – disease remission in patients with severe lupus (*15*). Nonetheless, excess morbidity and mortality from infection and cytokine-related toxicities observed with current immune effector cell therapies – including cytokine release syndrome (CRS) (*17*), immune effector cell-associated neurotoxicity syndrome (ICANS) (*18*), and delayed neutropenia (*19*), currently limit broad use, particularly for patients with less severe forms of SLE. These considerations motivate the development of precision immunotherapies that selectively eliminate autoreactive B cells while preserving protective immunity, thereby reducing the risk of infection. To date, analogous efforts have largely been focused on non-systemic autoimmune diseases that are organ-specific or driven by a single protein antigen (*20*). In these diseases, autoreactive B cells, though diverse, are distinguished from normal B cells by surface expression of BCRs that bind one self-antigen (*21*). This expression can be exploited through antigen-specific therapies that target these BCRs directly through their cognate autoantigen (*22–24*).

The development of antigen-specific therapeutic strategies for SLE is challenging in light of the extensive repertoire of autoantigens (>180) targeted by autoreactive B cells and their secreted autoantibodies (*25*), including nucleic acids (e.g., dsDNA) (*25–27*), proteins (e.g., histones, DNAse1L3, Sm, U1-ribonucleoprotein, Ro52, Ro60) (*28–30*), lipids (e.g., cardiolipin) (*28, 31*), and carbohydrates (e.g., I/i blood group antigens) (*32*). This diversity dramatically limits the feasibility of antigen-specific therapies in SLE and necessitates alternative strategies. One such approach, investigated in the current study, is to target autoreactive BCRs independently of their antigen-binding regions through unifying structural elements. In particular, the B cell compartment in SLE is uniquely characterized by a marked expansion of autoreactive B cell clones that use VH4-34 in their BCRs, irreversibly selected during V-D-J recombination in early B-cell development (*33*). The framework region 1 (FR1) of VH4-34 encodes the 9G4 idiotope (9G4id), a germline-encoded, conformational epitope that directly binds self-antigens (e.g., “i” and “I” blood group antigens and the glycoprotein CD45/B220) (*32, 34*). 9G4id B cells are therefore inherently autoreactive and normally restrained by peripheral tolerance checkpoints. In healthy individuals, they are excluded from germinal centers, and affinity-matured 9G4id B cells are therefore underrepresented in the memory B-cell compartment. However, in SLE, 9G4id B cells are licensed to participate in germinal center reactions, generating IgG+ memory B cells and plasma cells (*33*). These 9G4id B cells are sources of many of the autoantibodies in SLE (*28, 35, 36*), including antinuclear, anti-dsDNA, anti-histone, anti-DNAse1L3, anti-Sm/RNP, anti-Ro52, anti-Ro60, and anti-cardiolipin antibodies (autoreactive by virtue of binding of their affinity-matured complementarity determining regions [CDRs] in the context of germline-encoded FR1 of VH4-34), in addition to autoantibodies against red blood cell and immune cell antigens (autoreactive by virtue of binding through the germline-encoded FR1 of VH4-34 directly). 9G4 antibodies are specific for SLE (sensitivity 45-70%; specificity >95%) and correlate with disease activity (*37, 38*). Notably, SLE patients treated with belimumab, a BAFF-inhibiting monoclonal antibody, show a significant loss of VH4-34+ plasmablast (*39*). Additionally, lupus disease flares following peripheral B cell depletion with rituximab are characterized by rises in 9G4id anti-dsDNA antibodies (*40*). These data suggest that 9G4id B cells have a relatively rapid turnover after B cell-targeted therapies. Given their central role in active lupus and general dispensability in health, 9G4id B cells are a promising target for the depletion of autoreactive B cells in SLE.

In this study, we describe the invention of two precision cellular therapies targeting 9G4id B cells for the treatment of SLE and other 9G4id B cell-associated diseases.

## Results

### 9G4 cTCR-T cells and 9G4 CAR-T cells have similar capacity to bind the 9G4id

Pan-B cell-targeted cellular therapies that are currently being repurposed for the treatment of autoimmune diseases almost invariably use CARs as synthetic immune receptors. To design and comparatively test CAR-T cells expressing an anti-9G4 antibody (denoted “9G4-CAR”) for the selective targeting of 9G4id B cells, we grafted an anti-9G4 single-chain variable fragment (scFv), using the variable light (VL) chain-linker-variable heavy (VH) chain orientation, onto second-generation CAR scaffolds, using a CD28 costimulatory domain and a CD3ζ signaling domain in all constructs. We compared the performance of several modifications of this basic design, including those containing different linker peptides (a flexible poly-glycine-serine [G_4_S] or a more rigid five amino acid “EAAAK” linker), different hinge domains (CD8α, IgG4, CD3ζ, or [G4S]_3_), and different transmembrane domains (CD28 or CD3ζ). A total of five 9G4-CAR designs were tested with varying extracellular domain size, flexibility, and membrane anchoring to optimize immune synapse formation and to accommodate the large size of the target antigen (BCR) and the membrane-distal epitope “QW (15) AVY” (*36*) **(Fig. 1A, Fig. S1A)**.

**Figure 1.**
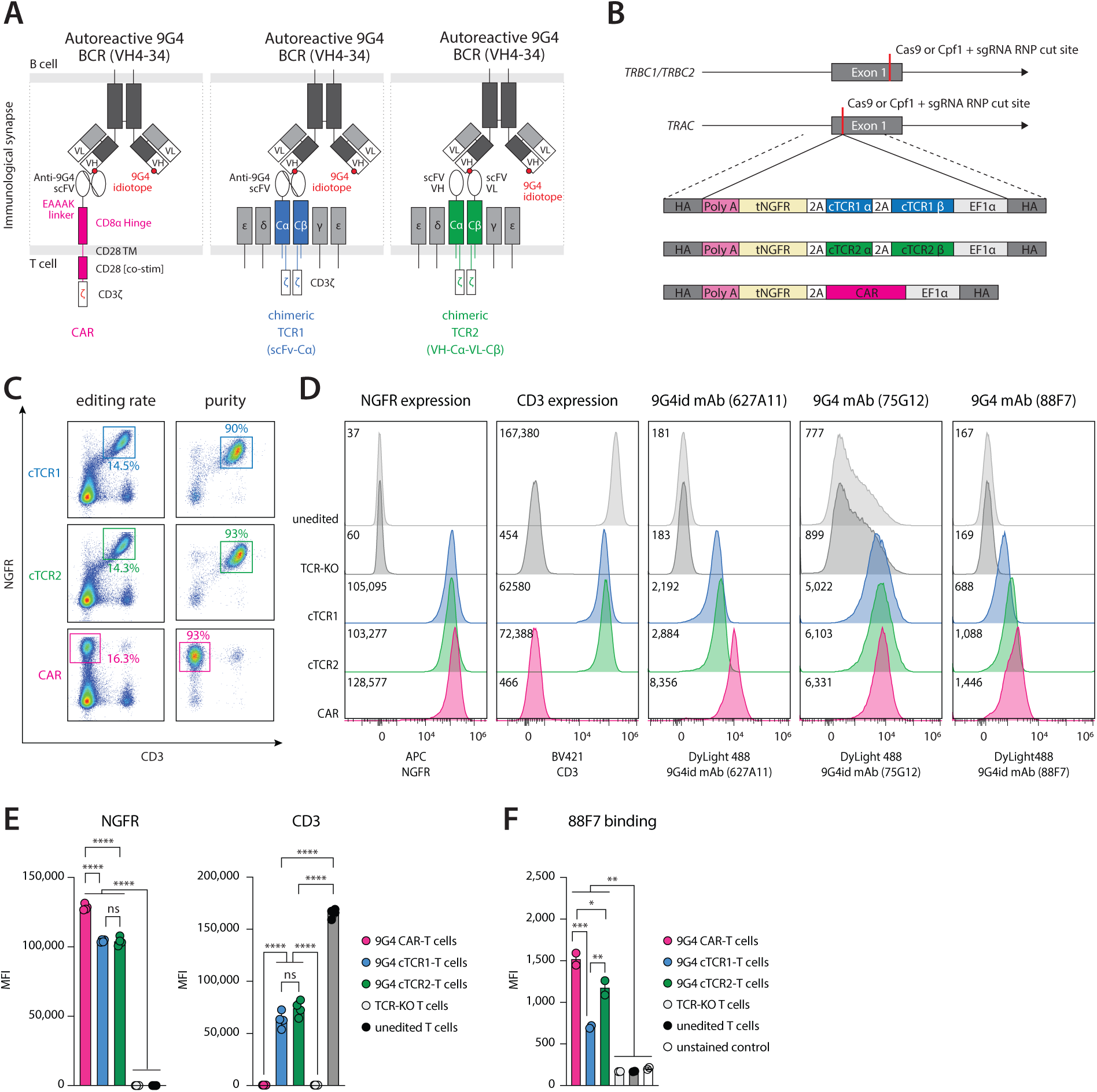
Expression of synthetic immune receptor in 9G4 CAR-T cells and 9G4 cTCR-T cells. **(A)** Diagrams of 9G4 cTCR1-, 9G4 cTCR2-T, and 9G4 CAR-T cells targeting 9G4id (VH4-34+) BCRs. The two cTCR constructs tested (cTCR1, cTCR2) are shown. cTCR1 comprises an anti-9G4 single-chain variable fragment (scFv) linked to a truncated TCR constant α (Cα) domain and a truncated TCR constant β (Cβ) domain, with the endogenous TCR variable domains removed. In comparison, cTCR2 comprises a split antibody fragment format that the immunoglobulin heavy (VH) chain of the anti-9G4 scFv to Cα and the immunoglobulin light (VL) chain to Cβ. TCR chains are similarly truncated to remove the endogenous TCR variable domains. The prioritized 9G4 CAR construct comprises an anti-9G4 scFv, an EAAAK linker, a CD8α hinge, a CD28 transmembrane (TM) domain, an intracellular CD28 co-stimulatory domain (co-stim), and an intracellular CD3ζ signaling domain. ε, δ, γ, and ζ denote CD3 ε, δ, γ and ζ subunits/domains, respectively. The autoreactive BCR containing the 9G4 idiotope (pink dot) is shown for comparison. **(B)** Visual summary of the CRISPR/Cas HDR strategy used to simultaneous knock out *TRBC1* and *TRBC2* (KO) and knock-in synthetic immune receptor coding sequences into *TRAC* (KI). cTCR1, cTCR2, and CAR homology directed repair templates (HDRTs) are shown specifically. Double-stranded DNA (dsDNA) HDRTs used comprise an EF1α promoter (EF1α) sequence, Kozak sequence, signal peptide sequence, the synthetic receptor domain(s), a tNGFR sequence (for selection), Stop codon, and a simian virus 40 polyadenylation (poly A) sequence. Different proteins are separated by furin-2A (2A) sequences for expression. Homology arms (HAs) for the target locus are approximately 300 base pairs (bps) in length, flanking the above sequences. **(C)** Flow cytometric validation of editing rates of engineered T cells four days after nucleofection (left column) and purity after positive bead selection (right column). Edited cells were visualized by staining for tNGFR (CAR, cTCR1, cTCR2 expected to be positive) and/or CD3 epsilon (cTCR1, cTCR2 expected to be positive). Percentage of edited single, live T cells is shown before and after positive selection. **(D)** Histograms (from left to right) of flow cytometric staining for tNGFR, CD3 epsilon (CD3), and three monoclonal 9G4id derived from SLE patients (clones 627A11, 75G12, and 88F7). Median Fluorescence Intensity (MFI) for each staining is shown in each panel. **(E)** Statistical summary of MFIs observed for tNGFR and CD3 for 9G4 CAR-T, 9G4 cTCR1-T, 9G4 cTCR2-T cells, TCR knock-out (KO)-T cells (control), and unedited T cells (control). Data are shown as mean ± SD of 4 replicates. **(F)** Statistical summary of MFIs observed for binding of monoclonal 9G4id 88F7 in the same set of T cells. Data are shown as mean ± SD of 2 replicates. \*\*\*\**P*<0.0001, \*\*\**P*<0.001, \*\**P*<0.01, \**P*<0.05, *P*=ns (not significant) by one-way ANOVA with Tukey’s multiple comparison test. All statistical analyses are provided in the Supplementary Dataset.

While CARs confer high cytotoxicity against cells that express target antigens at a high surface density (e.g., CD19), CAR-T cells have been shown to lack efficacy at low or even moderate surface antigen densities (<200-5000/cell) (*41, 42*). As B cells differentiate into plasma cells, alternative splicing skews immunoglobulin expression from membrane-bound to secreted forms, sharply reducing surface BCR density (*43, 44*). Precision cellular therapies that retain cytotoxicity at low or ultra-low antigen densities may thereby be more effective for autoimmune diseases where pathogenic B cells are also contained in the plasma cell compartment (i.e., diseases not appropriately treated by CD19 CAR-T cells). In contrast to CARs, natural T cells, using their TCRs, can mount effective cytotoxic responses against targets presenting very low antigen densities. Recent work has shown that re-engineered cTCRs retain TCR-like sensitivity and enable robust killing at very low antigen densities (<10 molecules per cell) (*45–47*). These designs fuse antibody fragments to a truncated TCR scaffold, thereby combining CAR-like binding with TCR-like signaling. To potentially extend the spectrum of targetable autoreactive B-cell lineage cells, we designed T cells that use re-engineered chimeric TCRs (cTCRs) as synthetic immune receptors **(Fig. 1A)**. For this purpose, we grafted anti-9G4 antibody fragments to the N-terminus of the TCRα and/or TCRβ chains, with endogenous variable domains removed, using two orthogonal designs **(Fig. 1A).** In one design, called 9G4-cTCR1, an anti-9G4 scFv was fused to a truncated TCRα chain via a rigid EAAAK linker, while the TCRβ variable domain (Vβ) chain was truncated. In a second design, called 9G4-cTCR2, the anti-9G4 scFv was split across the TCR: the VH domain was fused to truncated TCRα and the VL domain to truncated TCRβ, each joined by an EAAAK linker **(Fig. 1A)**.

To compare synthetic immune receptors under matched genomic and transcriptional contexts, we leveraged CRISPR/Cas (Cas9 or Cas12a/Cpf1)-mediated homology-directed repair (HDR) to introduce all constructs into the first exon of the TCRα constant (*TRAC*) gene locus of primary human T cells, simultaneously abrogating expression of the endogenous TCRα chain **(Fig. 1B)** (*47*). The TCRβ constant (*TRBC*) gene loci were concurrently knocked out to avoid mispairing and expression of the engineered T cells’ endogenous TCRs, a potential source of alloreactivity during extended co-culture killing assays. In all designs, expression of the cTCR or CAR transgene was driven by the elongation factor-1 alpha (EF1α) promoter, and a truncated nerve growth factor receptor (tNGFR) was included as a reporter for edited cell identification and purification (*47*). Controls included *TRAC*/*TRBC* double-knockout T cells (TCR-KO) and unedited T cells. Edited T-cell purity following CRISPR/Cas editing was measured by flow cytometry quantifying tNGFR and CD3 epsilon (CD3ε) expression, as each TCRαβ heterodimer associates with two CD3ε subunits (and CD3ε surface expression is lost after *TRAC*/*TRBC* KO without successful cTCR knock-in) (*48*). Initial editing efficiencies of approximately 15% were increased to 86-93% by positive selection for T cells expressing tNGFR (Methods, **Fig. 1C, Fig. S1B-C**).

Overall, tNGFR was expressed at comparable surface densities in all engineered anti-9G4 T cells **(Fig. 1C-E, Fig. S1B-E)**, confirming successful editing of *TRAC*. CAR designs showed comparable surface tNGFR expression among each other (**Fig. S1D-E**), and modestly higher expression compared to cTCR designs **(*P*<0.0001; Fig. 1D-E)**, possibly due to the smaller CAR constructs’ size. As expected, 9G4-CARs and TCR-KO T cells had no detectable CD3ε surface expression as a result of the disruption of TCRα and TCRβ constant chains, which preclude surface trafficking of CD3ε **(Fig. S1D-E)**. By contrast, 9G4-cTCR constructs (cTCR1, cTCR2) showed only a 2-fold lower surface CD3ε expression than unedited T cells **(*P*<0.0001; Fig. 1D-E)**.

To directly compare the expression and functional availability of the relevant antibody fragments in all synthetic immune receptors, we stained engineered anti-9G4 T cells with established monoclonal 9G4id antibodies (clones 627A11, 75G12, 88F7) (*28*). Despite comparable tNGFR expression across CAR designs, the 9G4 CAR with EAAAK-CD8α-Hinge domains showed higher binding capacity to 9G4id than other CARs **(*P*<0.0001; Fig. S1D-F)**, and higher binding to 9G4id compared to 9G4-cTCR constructs **(*P*<0.05)**. cTCR2-T cells bound slightly more to monoclonal 9G4id than cTCR1-T cells **(*P*<0.01; Fig. 1D-F)**. Interestingly, antibody clone 75G12 stained TCR-KO T cells and unedited T cells independent of anti-9G4 synthetic immune receptor expression **(Fig. 1D, Fig. S1D)**, mirroring its reported ability to non-specifically bind other immune cells (*36*). Together, these results confirmed the successful engineering of 9G4-targeted synthetic immune receptors in primary human T cells and revealed design--specific differences in receptor display and binding.

### 9G4 CAR-T cells and 9G4 cTCR-T cells selectively kill SLE-9G4id B cell lines with comparable potency

To functionally test the potency and specificity of these engineered T cells, we generated model B cell lines by replacing the native BCR of Ramos B cells, a human Burkitt lymphoma cell line, with monoclonal BCRs of interest. This was achieved by editing the *IGH* locus of Ramos B cells using CRISPR/Cas9 HDR to express a single-chain BCR/antibody construct containing a heavy chain constant region splice junction (*49*). This process resulted in expression of both membrane-bound (BCR) and secreted forms (antibody) of the transgenic monoclonal immunoglobulin under the control of endogenous regulatory elements. All BCR/antibody constructs were designed as a single-chain molecule linking the desired light chains and variable heavy chains with a Strep-tag II-containing linker peptide, enabling detection and positive selection of surface-expressed BCRs **(Fig. 2A-B)**. Using this approach, we generated autoreactive B cell lines expressing one of three SLE patient-derived 9G4id monoclonal BCR/antibodies (SLE 9G4id BCR_627A11_, BCR_75G12_, and BCR_88F7_, respectively) and a control B cell line expressing a non-9G4id (VH4-4+) monoclonal BCR/antibody (BCR_VH4-4_) **(Fig. 2C)**.

**Figure 2.**
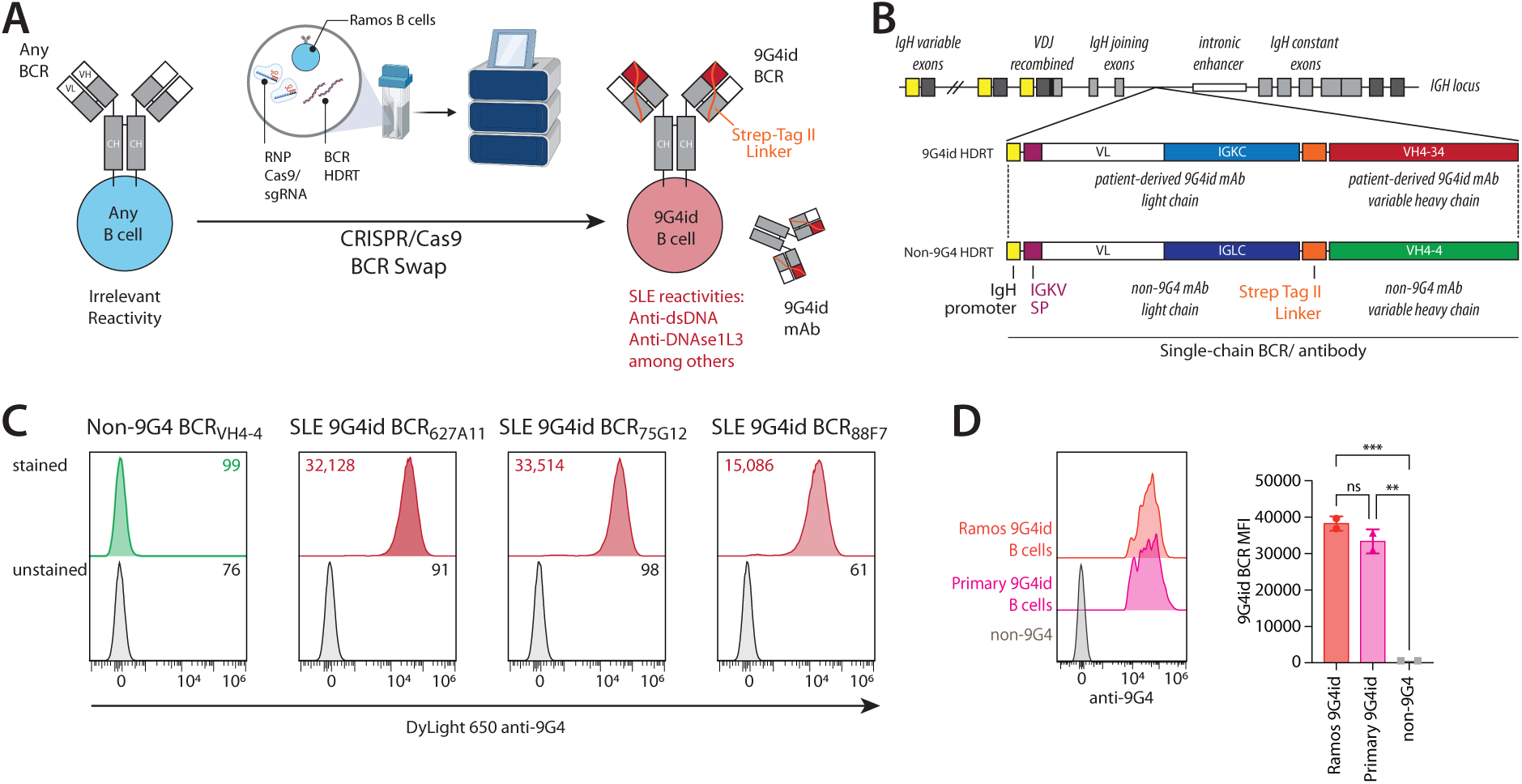
Generation of autoreactive Ramos B cell lines expressing SLE patient-derived, 9G4id BCRs/ antibodies. **(A)** Schematic of the experimental strategy to replace the endogenous BCR/antibody of Ramos B cells with desired patient-derived autoreactive BCR/ secreted antibody using CRISPR/Cas9-mediated HDR. This strategy was employed to generate three Ramos B cell lines expressing autoreactive 9G4id (VH4-34+) BCRs from patients with SLE and one control BCR (VH4-4+). CRISPR, Clustered Regularly Interspaced Short Palindromic Repeats; RNP, ribonucleoprotein; sgRNA, single-guide RNA; HDRT, homology-directed repair template; dsDNA, double-stranded DNA. **(B)** Visual summary of the CRISPR/Cas9 HDR gene editing strategy used to introduce the paired immunoglobulin variable light (VL) and variable heavy (VH) chains of monoclonal BCR/antibody into the *IGH* gene locus, abrogating expression of the endogenous BCR. Monoclonal immunoglobulin of interest is expressed as a single-chain construct, incorporating a Strep-tag II linker used for detection and positive selection. *IGKC*, immunoglobulin kappa constant chain; *IGLC*, immunoglobulin lambda constant chain; *IGKV*, immunoglobulin kappa variable chain; SP, signal peptide. **(C)** Flow cytometric staining of engineered Ramos B cell clones (non-9G4 [VH4-4] B cell clone, left panel; 9G4id SLE B cell clones, right panels) with an anti-9G4 antibody, after magnetic bead enrichment and single clone selection. Median Fluorescence Intensity (MFI) of each stained and unstained B cell clone is shown. **(D)** Comparison of 9G4id BCR expression levels on engineered SLE 9G4id Ramos BCR627A11 cells and primary human B cells by flow cytometric staining with an anti-9G4 antibody. Non-9G4 B cells are shown as a negative control. Data are shown as mean ± SD of two technical replicates. Data are representative of n = 2 independent experiments. \*\*\**P*<0.001, \*\**P*<0.01, *P*=ns (not significant) by one-way ANOVA with Tukey’s multiple comparison test. All statistical analyses are provided in the Supplementary Dataset.

Surface expression of 9G4id BCRs on autoreactive Ramos B cells (SLE 9G4id BCR_627A11_, BCR_75G12_, and BCR_88F7_) and absence of 9G4id BCRs on non-9G4 control Ramos B cells (BCR_VH4-4_) were confirmed by flow cytometry using a monoclonal anti-9G4 antibody **(Fig. 2C)**. We also determined the surface expression levels of 9G4id BCRs on engineered 9G4id Ramos B cells and primary human 9G4id B cells isolated from human PBMCs by flow cytometry. BCR densities on engineered 9G4id B cells were similar to the surface expression levels on primary human 9G4id B cells **(*P*=0.22; Fig. 2D)**.

We next compared the potency and specificity of our engineered immune effector cells in co-culture with one of these B-cell models (SLE 9G4id Ramos BCR_672A11_ B cells). Compared to TCR-KO control T cells, which did not deplete target B cells or control B cells, all designs of 9G4 CAR-T cells showed complete elimination of SLE 9G4id BCR_627A11_ B cells, while preserving non-9G4 BCR_VH4-4_ Ramos B cells (control vs CAR-T, *P*<0.0001 for all effector-to-target cell [E:T] ratios) **(Fig. S1G)**. While no major differences in cytotoxicity were observed among the tested 9G4 CAR-T cell designs **(Fig. S1G)**, we prioritized the construct incorporating an EAAAK-CD8α hinge (hereinafter “9G4 CAR-T cells”) based on its superior receptor surface expression (**Fig. S1D-F**). This 9G4 CAR-T cell construct was therefore used for subsequent comparisons with 9G4 cTCR1-T and cTCR2-T cells.

We next comparatively evaluated the potency and specificity of 9G4 CAR-T cells, 9G4 cTCR1-T cells, and 9G4 cTCR2-T cells using our Ramos B cell model cell lines. We performed cytotoxicity assays in which we co-cultured 9G4 cTCR-T cells or 9G4 CAR-T cells with one of three SLE 9G4id Ramos B cell lines (BCR_672A11_, BCR_75G12_, or BCR_88F7_) or with non-9G4 Ramos B cells (BCR_VH4-4_). After four days of co-culture, we quantified the absolute number of live, single Ramos B cells (GFP+) by flow cytometric staining of StrepTag II presented in edited BCRs **(Fig. 3A)**. Compared to control T cells, co-culture with 9G4 CAR-T cells, 9G4 cTCR1-T cells, and cTCR2-T cells led to a marked reduction in all 9G4id Ramos B cells (*P<*0.05 for all E:T ratios tested) **(Fig. 3B)**, while non-9G4 Ramos B cells were not depleted. Next, we compared the cytotoxicity derived from different T cells by live-cell imaging, longitudinally quantifying Ramos B cells by their expression of green fluorescent protein (GFP) **(Fig. 3C)**. In a four-day co-cultures, 9G4 CAR-T cells, 9G4 cTCR1-T cells, and 9G4 cTCR2-T cells all controlled target 9G4id Ramos cell growth (*P*<0.0001 compared to control T cells) **(Fig. 3D-G, Video 1)**. In contrast, non-9G4 Ramos B cell growth was not significantly impacted by co-culture with anti-9G4 T cells **(*P>*0.23; Fig. 3C-D, 3H, Video 2)**. These results also showed no significant differences in cytotoxicity between 9G4 CAR-T cells and 9G4 cTCR-T cells **(*P>*0.90; Fig. 3D)**.

**Figure 3.**
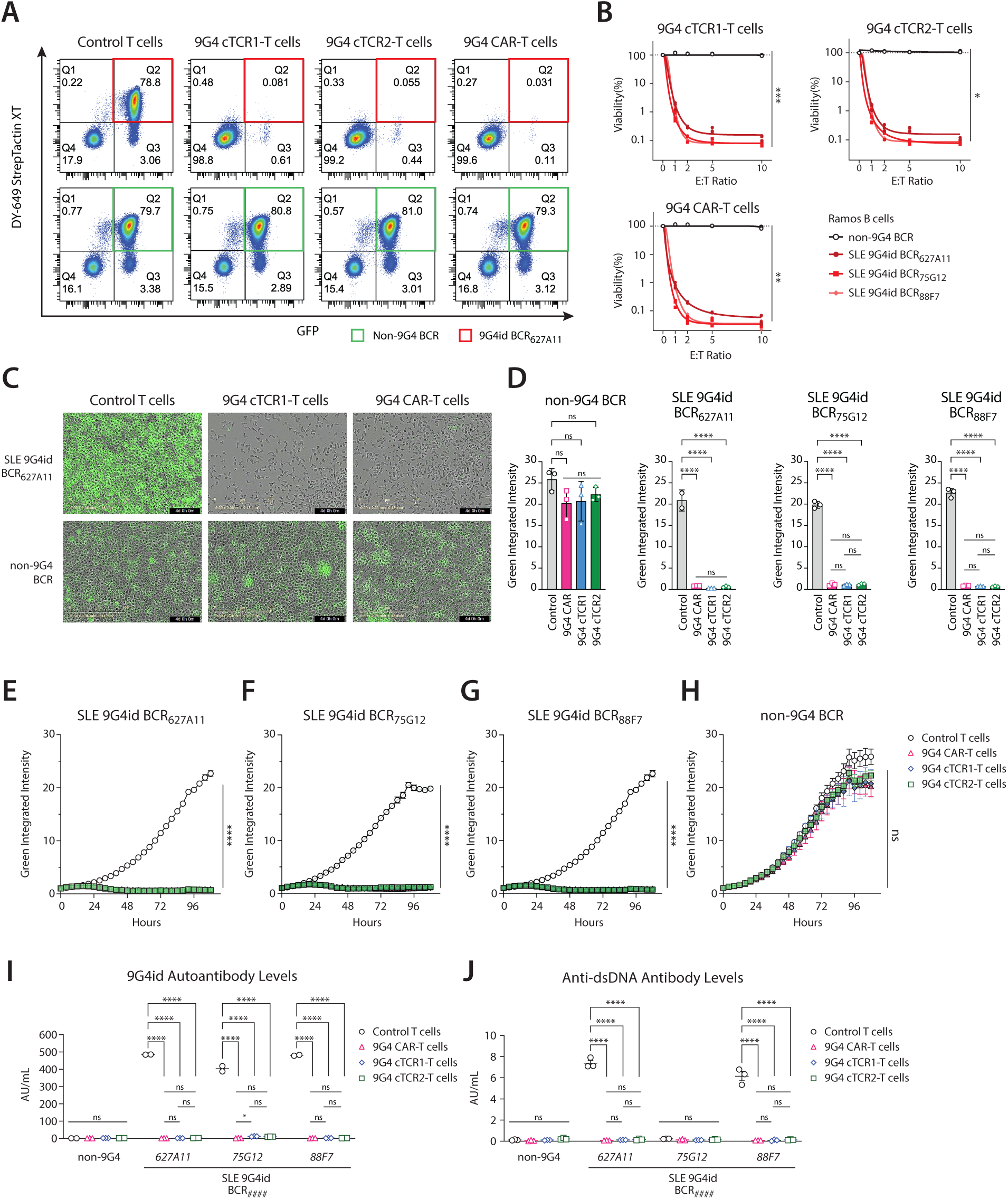
T cells engineered to express anti-9G4 synthetic immune receptors eliminate 9G4id B cells with comparable specificity and potency. **(A)** Representative flow cytometry plots illustrating selective killing of Ramos B cells expressing the SLE patient-derived, autoreactive 9G4id BCR_627A11_ (top row) at the end of co-culture with either control T cells, 9G4 cTCR1-T cells, 9G4 cTCR2-T cells, or 9G4 CAR-T cells (top panels, from left to right). 9G4id BCR_627A11_ B cells (Q2, top row), known to react with lupus antigens dsDNA, DNAse1L3, and cardiolipin, were eliminated by 9G4 cTCR1-T cells, 9G4 cTCR2-T cells, and 9G4 CAR-T cells, but not control T cells. Ramos B cells expressing a non-9G4 BCR/antibody (VH4-4), used as an irrelevant B cell control, were not depleted under the same conditions (Q2, bottom row). In this example, flow cytometric analysis of single, live cells after 4 days of co-incubation at an effector-to-target cell [E:T] ratio of 10:1 (100,000 edited T cells, 10,000 GFP+ Ramos B cells) is shown. B cells are identified as GFP+, DY649 StrepTactin XT+ cells (binding Strep-tag II in surface-expressed BCRs, Q2). T cells are visualized in Q4 as GFP-, StrepTactin XT- cells. **(B)** Percent viability of Ramos B cells (SLE 9G4id Ramos BCR_627A11_, SLE 9G4id Ramos BCR_75G12_, SLE 9G4id Ramos BCR_88F7_, or non-9G4 BCR) at the end of co-culture with either 9G4 cTCR1-T cells (top left graph), 9G4 cTCR2-T cells (top right graph), or 9G4 CAR-T cells (bottom left graph) for different E:T ratios (E:T=10:1, 100,000 edited T cells; E:T=5:1, 50,000 edited T cells; E:T=2.5:1, 25,000 edited T cells; E:T=1.25:1, 12,500 edited T cells; 10,000 B cells). The absolute numbers of single, live GFP+, StrepTactin XT+ B cells, as determined by flow cytometry, were used to calculate % viability. All data were normalized to the co-culture condition with control T cells (E:T=0). \*\*\**P*<0.001, \*\**P*<0.01, \**P*<0.05; two-way ANOVA with Dunnett’s multiple comparison test. **(C-H)** Co-culture of GFP+ Ramos B cells (non-9G4 BCR, BCR_627A11_, BCR_75G12_, or BCR_88F7_) with engineered anti-9G4 T cells (9G4 CAR, 9G4 cTCR1, or 9G4 cTCR2) or control T cells at E:T 5:1, longitudinally analyzed by live-cell imaging for 108 hours (Incucyte SX5, 20x). **(C)** Representative images showing GFP+ Ramos B cells (non-9G4 BCR, bottom row; 9G4id BCR_627A11_, top row) in co-culture with control T cells (left panels), 9G4 cTCR1-T cells (middle panels), or 9G4 CAR-T cells (right panels) at 4 days (20x, phase contrast and green channel). **(D)** Quantification of Green Integrated Intensity of GFP+ Ramos B cells (panels from left to right: non-9G4 BCR, BCR_627A11_, BCR_75G12_, or BCR_88F7_) in co-culture with T cells (control T cells [Control], 9G4 CAR, 9G4 cTCR1, or 9G4 cTCR2). All data were normalized to day 0 of co-culture in the same condition. \*\*\*\**P*<0.0001, ns, not significant; one-way ANOVA with Tukey’s multiple comparison test. **(E-H)** Changes in Green Integrated Intensity in co-cultures with GFP+ Ramos BCR_627A11_ (E), BCR_75G12_ (F), BCR_88F7_ (G), or non-9G4 BCR B cells (H), as quantified by live-cell imaging. **(I)** Quantification of secreted 9G4id antibody levels by ELISA in culture supernatants of non-9G4 BCR Ramos B cells, BCR_627A11_ Ramos B cells, BCR_75G12_ Ramos B cells, or BCR_88F7_ Ramos B cells incubated with control T cells or engineered anti-9G4 T cells (9G4 CAR, 9G4 cTCR1, or 9G4 cTCR2). Comparisons are representative of n=2 experiments. **(J)** Quantification of secreted anti-dsDNA autoantibody levels by ELISA in the same co-culture supernatants. Comparisons are representative of n=2 experiments. Data are shown as mean ± SD of technical replicates. \*\*\*\**P*<0.0001, \**P*<0.05, ns, not significant; two-way ANOVA with Tukey’s multiple comparison test. All statistical analyses are provided in the Supplementary Dataset.

Repeated stimulation assays (RSAs) can showcase early T cell exhaustion and have been shown to be good predictors of in-vivo efficacy with other immunotherapeutic approaches (*47*). We therefore extended our model by administering additional B cells every 3 days to a 12-day co-culture of engineered anti-9G4 T cells and Ramos B cells. As before, we observed that 9G4 CAR-T cells, 9G4 cTCR1-T cells, and 9G4 cTCR2-T cells maintained robust and comparable potency against SLE 9G4id Ramos B cells through at least one additional challenge, without impairing the growth of non-9G4 B cells **(Fig. S2A-D)**. With a second challenge, 9G4 CAR-T cells, 9G4 cTCR1-T cells, and 9G4 cTCR2-T cells maintained strong and comparable potency for two SLE 9G4id Ramos B cell lines (BCR_627A11_ and BCR_88F7_), while cytotoxicity appeared to decline against SLE 9G4id BCR_75G12_ Ramos B cells **(Fig S2A-C)**. Interestingly, this outgrowth in SLE 9G4id BCR_75G12_ co-culture conditions was driven by BCR_75G12_-negative B cells and was more pronounced with cTCR-T cells than with CAR-T cells **(cTCR1/cTCR2 vs CAR, *P*<0.01; Fig. S2B-D)**. These data suggest antigen escape in this engineered malignant Ramos B cell line, which is not expected when targeting primary B cells in patients with autoimmune disease. Co-culture with 9G4 CAR-T cells resulted in reduced expansion of BCR_75G12_-negative populations (CAR vs control T cells, *P*<0.001) **(Fig. S2B)**, suggesting higher fitness or increased cytokine-mediated unspecific cytotoxicity.

We assumed that the depletion of autoreactive 9G4id B cells would similarly impair autoantibody production. To validate this assumption, we measured 9G4id antibody levels and anti-dsDNA antibody levels in the co-culture supernatants by Enzyme-Linked Immunosorbent Assay (ELISA). Compared to control T cells, we observed a consistent reduction in 9G4id antibody production after co-culture with engineered anti-9G4 T cells (control vs CAR-T, cTCR1, and cTCR2, respectively, *P*<0.0001) **(Fig. 3I)**. Similar reductions of anti-dsDNA antibodies were observed in experiments with B cells expressing 9G4id antibody clones 627A11 and 88F7 **(*P*<0.0001; Fig. 3J)**. The 9G4id antibody clone 75G12 is known to not bind to dsDNA and served as a control in this experiment.

### Engineered anti-9G4 T cells deplete 9G4id B cells in cold agglutinin disease and clonal B cell disorders

We next investigated whether engineered anti-9G4 T cells had similar efficacy against 9G4id B cells that drive other disease states. Cold agglutinins (CAs) are cold-activated antibodies that bind to red blood cells at low temperature, resulting in their agglutination and autoimmune hemolytic anemia (*50, 51*). In patients with cold agglutinin disease (CAD), a clonal B-cell lymphoproliferative disorder, IgM CAs are almost exclusively encoded by the VH4-34 gene and specific for I/i antigens expressed on red blood cells (bound by the 9G4id) (*52–56*). To confirm whether engineered anti-9G4 T cells could eliminate B cells driving CAD, we generated three 9G4id Ramos B cell lines expressing different CAD patient-derived, CA+ BCRs (CAD 9G4id BCR_KAU_, BCR_FS-2_, and BCR_FS-1_, respectively). Surface expression levels of the 9G4id BCRs were confirmed using monoclonal anti-9G4 antibodies by flow cytometry **(Fig. S3A)**. To evaluate the specificity and cytotoxicity of 9G4 CAR-T cells and 9G4 cTCR1-T cells against CAD 9G4id B cells, we co-cultured CAD 9G4id Ramos B cells with engineered anti-9G4 T cells or control T cells **(Fig. S3B)**. All CAD 9G4id B cell clones were eliminated by both 9G4 CAR-T cells and 9G4 cTCR1-T cells (*P*<0.01 at all E:T ratios tested) **(Fig. S3C-D),** while non-9G4 BCR_VH4-4_ cells were preserved. Interferon (IFN)-γ secretion by anti-9G4 synthetic immune receptor T cells was detected only in the presence of target cells, as quantified in co-culture supernatants by ELISA **(Fig. S3E)**.

Over-representation of the 9G4id has been reported in several types of B cell cancers (*57–61*). To confirm that our engineered anti-9G4 T cells could recognize and kill such cancer cells, we used the experimental paradigm described above but with wild-type Ramos (RA1) B cells as the target cells. Ramos (RA1) cells naturally express endogenous (not engineered) VH4-34 and are a canonical Burkitt lymphoma cell line. After confirming surface expression of endogenous 9G4id BCR **(Fig. S3F)**, we co-incubated 9G4 CAR-T cells and 9G4 cTCR1-T with wild-type Ramos B cells, resulting in the efficient elimination of target B cells **(Fig. S3G-H)**. IFN-γ was secreted by engineered anti-9G4 T cells only in the presence of target cells and in a dose-dependent fashion **(Fig. S3I).**

### 9G4 cTCR-T cells evoke lower cytokine release than 9G4 CAR-T cells in the treatment of autoimmune diseases

Cytokine-related toxicities, including CRS and ICANS, are known and potentially life-threatening complication of immune effector cell therapy, commonly observed in patients (*17, 18*). While these toxicities tend to be less severe in patients with autoimmune diseases, cytokine-related toxicities remain a major obstacle to more widespread adoption of these therapies. Strategies that can reduce these risks may allow cellular therapies to be applied to patients with less severe disease and in settings that require less intense monitoring. To understand how the use of different synthetic immune receptors may modulate the risk of cytokine-related toxicities, we quantified cytokine release and T-cell activation in co-cultures. Overall, cytokine secretion did not correlate with improved cytotoxicity in our study. For example, 9G4 CAR-T cells, when co-cultured with 9G4id B cells, secreted significantly higher levels of cytokines than 9G4 cTCR1-T cells or 9G4 cTCR2-T cells **(Fig. S3E, S3I; Fig. 4A-B; Fig. S4A-I)**, despite comparable effectiveness in killing target B cells. Specifically, 9G4 cTCR1-T cells and 9G4 cTCR2-T cells released significantly lower levels of IFN-γ than 9G4 CAR-T cells **(*P*<0.01; Fig. 4A)**. Granzyme A and B (*P*<0.0001), IL-2Rα (*P*<0.0001), IL-4 (*P*<0.01), IL-6 (*P*<0.0001), IL-8 (*P*<0.0001), TNF-α (*P*<0.0001), TNF-β (*P*<0.0001), and GM-CSF (*P*<0.0001) were similarly reduced in the 9G4 cTCR-cells compared to the 9G4 CAR-T cells **(Fig. 4B)**.

**Figure 4.**
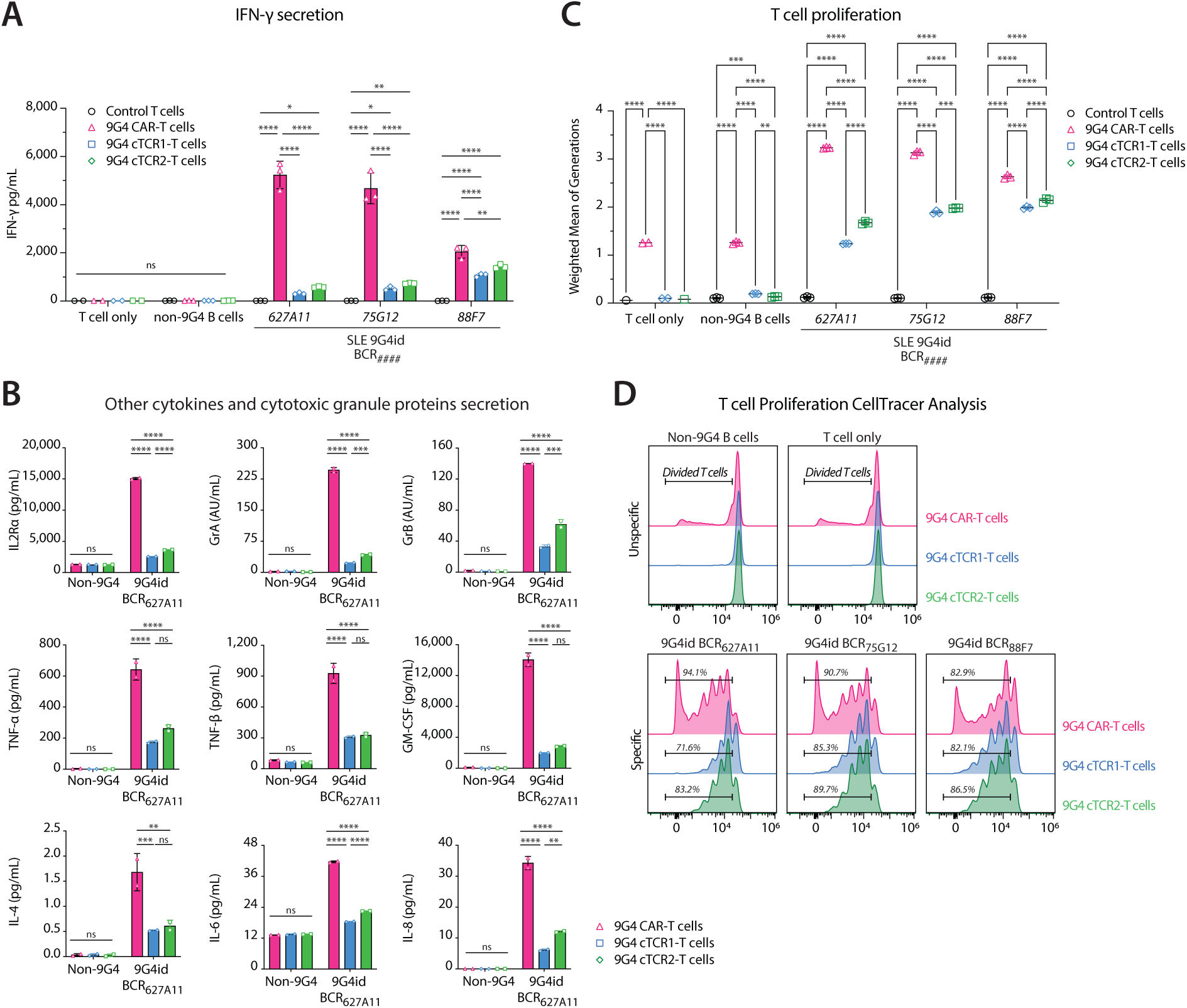
9G4 cTCR-T cells restrain cytokine release and prevent antigen-independent expansion. **(A)** To allow for direct comparison, different 9G4 synthetic immune receptor T cells (9G4 CAR-T cells, 9G4 cTCR1-T cells, 9G4 cTCR2-T cells) or control T cells were cultured with the SLE 9G4id Ramos BCR_627A11_, BCR_75G12_, BCR_88F7_, or non-9G4 B cells (E:T=5:1, 62 hours). Quantification of IFN-γ in conditioned co-culture supernatants by ELISA is shown. Columns and error bars represent mean ± SD. **(B)** Quantification of other cytokines (IL2Rα, TNF-α, TNF-β, GM-CSF, IL-4, IL-6, IL-8) and secreted cytotoxic granule proteins (granzyme A [GrA], granzyme B [GrB]) in co-culture supernatants of the same experiment shown in A (U-PLEX assay, Meso Scale). Columns and error bars show mean ± SD. **(C)** Proliferation of T cells (9G4 CAR-T cells, 9G4 cTCR1-T cells, 9G4 cTCR2-T cells, or control T cells) in response to target B cells (SLE 9G4id Ramos BCR_627A11_, BCR_75G12_, BCR_88F7_) or irrelevant B cells (non-9G4 BCR Ramos B cells) as determined by fluorescent dye dilution. Anti-9G4 synthetic immune receptor T cells and control T cells labeled with CellTrace Violet (CTV) were cultured without B cells (“T cell only”), with non-9G4 BCR Ramos B cells, or with SLE 9G4id Ramos B cells (BCR_627A11_, BCR_75G12_, BCR_88F7_) at an E:T ratio of 5:1. Flow cytometry was performed after 108 hours of co-incubation, and CTV dye dilution quantified in live, single T cells. Weighted average of generations (mean ± SD) was calculated to quantify T-cell proliferation for statistical analysis. **(D)** Flow cytometric analysis of CTV dilution in single, live T cells, showing the percent of divided, engineered T cells (9G4 CAR, 9G4 cTCR1, or 9G4 cTCR2) in the presence of irrelevant B cells (non-9G4 B cells, top left panel), no B cells (“T cells only”, top right panel), or SLE 9G4id Ramos B cell lines (bottom row panels). Comparisons are representative of n=2 independent experiments. \*\*\*\**P*<0.0001, \*\*\**P*<0.001, \*\**P*<0.01, \**P*<0.05, ns=not significant; two-way ANOVA with Tukey’s multiple comparison test.

The increased cytokine release by 9G4 CAR-T cells was associated with increased T-cell proliferation following co-culture with 9G4id B cells (CAR-T vs cTCR1 and cTCR2, coincubation with all 9G4id B cells, *P*<0.0001) **(Fig 4C-D)**. Notably, proliferation of T cells independent of 9G4id B cells was observed for 9G4 CAR-T cells but did not occur in 9G4 cTCR-T cells **(*P*<0.0001; Fig 4C-D)**.

### Engineered anti-9G4 T cells selectively deplete autoreactive B cells in patients with SLE

We next tested the potency and specificity of anti-9G4 T cells against primary human B cells isolated from patients with SLE **(Table S1)**. The frequencies of 9G4id peripheral blood B cells in SLE patients’ PBMCs were determined by flow cytometric staining with anti-CD19 and anti-9G4 antibodies. Total B cells comprised 7.31% (patient 1), 5.99% (patient 2), 5.3% (patient 3), and 3.78% (patient 4) of total PBMCs. 9G4id B cells comprised 6.3% (patient 1), 5% (patient 2), 5% (patient 3), and 6.11% (patient 4) of total B cells **(Table S2)**. Co-cultures were conducted using PBMCs and donor-matched T cells to eliminate alloreactivity. For this experiment, primary human T cells were isolated from PBMCs of selected SLE patients, activated, and engineered to generate autologous 9G4 CAR-T cells, 9G4 cTCR-T cells, and second-generation CD19 CAR-T cells, using the same strategy described above for editing primary human T cells from healthy individuals (editing rate, purity after enrichment, and T-cell functional properties are summarized in **Table S2**).

Edited or unedited T cells were then co-cultured with matched patient PBMCs. After a 6-day co-culture in B-cell stimulation media, cells were transferred to FluoroSpot plates and incubated in the same media for another 24 hours to further promote B cell differentiation into antibody-secreting cells (ASCs). Subsequently, the number of 9G4id IgG ASCs and total IgG ASCs were quantified using a FluoroSpot assay **(Fig. 5A)**. Incubation with 9G4 CAR-T cells and 9G4 cTCR-T cells resulted in 95-100% reduction in the number of IgG+ 9G4id ASCs compared to control T cells **(*P*<0.0001; Fig. 5B**), without significant reduction of total IgG+ ASCs (*P*=0.90) **(Fig. S5A)**. By contrast, autologous CD19 CAR-T cells depleted 100% of IgG+ ASCs compared to control T cells **(Fig. S5A)**. We further confirmed the FluoroSpot results by bulk mRNA BCR repertoire sequencing **(Fig. 5C-D, Fig. S5F-G)**. After co-culture of PBMCs with either 9G4 CAR-T cells or 9G4 cTCR-T cells, over 99.7% of the 9G4id B-cell compartment was depleted compared to control T cells (*P*<0.05) **(Fig. 5D)**, while other B cell compartments were not depleted (*P*>0.05 for various IHGV genes). We observed that B cells expressing IGHV2-26, IGHV6-1, and IGH4-30-2 genes expanded after co-incubation, and no other compartments were depleted. To further validate these findings, we cocultured 9G4 CAR-T or 9G4 cTCR-T cells with primary B cells from healthy donors. Flow cytometric staining with anti-9G4 antibody, which detects all isotypes (IgG, IgA, IgM) of 9G4id B cells **(Fig. S5B)**, confirmed selective depletion of 9G4id B cells (*P*<0.05) **(Fig. S5D),** while preserving the overall B-cell compartment (*P*>0.1) **(Fig. S5C)**. At relatively lower E:T ratios (0.3:1 and 0.6:1), 9G4 CAR-T cells demonstrated approximately two-fold greater cytotoxicity than 9G4 cTCR-T cells (*P*<0.05) **(Fig. S5D)**, but secreted significantly higher levels of IFN-γ (1315 pg/mL vs. undetectable in cTCR1-T cells, *P*<0.0001 at E:T = 0.6:1) **(Fig. S5E)**.

**Figure 5.**
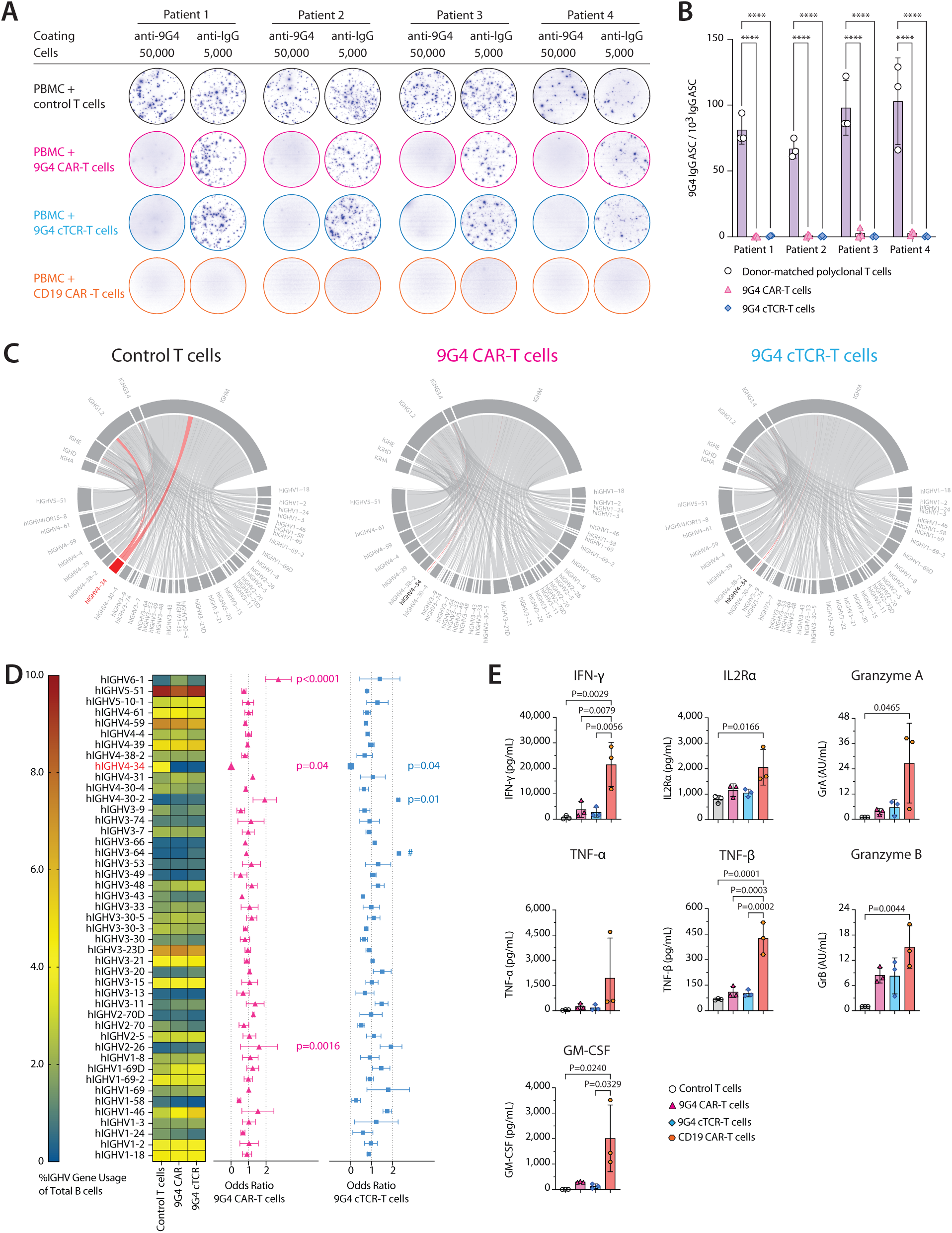
9G4 CAR-T cells and 9G4 cTCR-T cells selectively eliminate 9G4id B cells in SLE patient PBMCs. **(A)** Results of B cell FluoroSpot assays showing selective killing of primary human 9G4id B cells/antibody secreting cells (ASCs) from patients with SLE in co-culture with autologous 9G4 CAR- or 9G4 cTCR-T cells. For this experiment, 100,000 PBMCs from SLE patients (n=4) were co-cultured with 5,000 9G4 donor-matched control T cells (autologous CD3+ T cells, no genetic modification), 9G4 CAR-T cells, 9G4 cTCR1-T cells, or CD19 CAR-T cells in B-cell stimulation/ differentiation media. After 6 days, cells were transferred to FluoroSpot 96-well plates coated with either anti-9G4 antibody (50,000 cells/ well) or anti-human IgG antibody (anti-hIgG, 5,000 cells/well). After 20 hours of incubation, spots corresponding to 9G4id+ ASCs or total hIgG+ ASCs were detected using fluorophore conjugated anti-hIgG antibody (shown in pseudocolor). **(B)** Quantification of results shown in A. The number of IgG 9G4id ASCs per 1,000 total IgG ASCs is shown. Autologous 9G4 CAR-T cells and 9G4 cTCR1-T cells both eliminated IgG 9G4id ASCs from patients with SLE. Two-way ANOVA with Tukey’s multiple comparison test. **(C)** IGHV mRNA sequencing was performed at the end of co-culture of SLE PBMCs with either autologous control T cells (left), 9G4 CAR-T cells (middle), or 9G4 cTCR1-T cells (right). Chord diagrams of IGHV and IGHC gene usage show depletion of IGHV4-34+ B cell clonotypes which are predominantly found in the IgG1/2 and IgM compartments. Data shown for patient 3 as a presentative example. **(D)** Left, Heat map summarizing the usage of different IGHV genes (in % of total B cells) of B cells from SLE patients 1-3 after co-incubation with control T cells, 9G4 CAR-T cells, or 9G4 cTCR1-T cells, as determined by bulk repertoire sequencing. Right, forest plots depicting mean odds ratios and SD for IGHV gene usage in PBMCs (n=3) treated with donor-matched control T cells vs 9G4 CAR-T cells (pink tringles, left) or 9G4 cTCR1-T cells (blue squares, right). #denotes only one sample. One-way ANOVA with Dunnett’s multiple comparison. **(E)** Quantification of cytokines (IFN-γ, IL2Rα, TNF-α, TNF-β, GM-CSF) and secreted cytotoxic granule proteins (granzyme A [GrA], granzyme B [GrB]) in co-culture supernatants of SLE patient PBMCs (n=3) treated with 9G4 CAR-T cells, 9G4 cTCR1-T cells, conventional CD19 CAR-T cells, or control T cells at 48 hours (U-PLEX assay, Meso Scale). Data are shown as mean ± SD. \*\*\*\**P*< 0.0001, \*\*\**P*<0.001, \*\**P*<0.01, \**P*<0.05; one-way ANOVA with Tukey’s multiple comparison.

To evaluate potential differences between the anti-9G4 T cells described here and previously described CD19 CAR-T cells, we interrogated cytokine secretion at 48 hours of co-incubation of engineered or control T cells with SLE patient PBMCs. Compared to autologous CD19 CAR-T cells, both 9G4 CAR-T cells and 9G4 cTCR-T cells tended to produce lower levels of effector cytokines and proteases, including IFN-γ (*P*=0.003), IL2Rα (*P*=0.025), Granzyme A (*P*=0.056), and Granzyme B (*P*=0.079) **(Fig. 5E)**. Additionally, CD19 CAR-T cells produced higher amounts of pro-inflammatory cytokines that are commonly associated with CRS(*62*), such as TNF-β (*P*<0.0001) and GM-CSF (*P*=0.024) **(Fig 5E).**

### 9G4 CAR-T cells and 9G4 cTCR-T cells selectively eliminate autoreactive 9G4id B cells in vivo

To evaluate the efficacy of engineered anti-9G4 T cells in vivo, we established a humanized xenograft model engrafting autoreactive SLE 9G4id Ramos B cells into NSG mice. This model allowed us to test our therapeutic candidates in the context of B cells that express dsDNA/DNase1L3-reactive BCRs and secrete IgM anti-dsDNA/ anti-DNase1L3 antibodies that might bind and neutralize anti-9G4 T cells. Mice were injected with bioluminescent SLE 9G4id Ramos BCR_627A11_ B cells, followed by treatment with either 9G4 CAR-T cells, 9G4 cTCR-T cells, or control (TCR KO) T cells two days later **(Fig. 6A)**. Mice maintained stable body weights throughout the experiment **(Fig. 6B)**. Serial bioluminescence imaging (BLI) demonstrated robust expansion of 9G4id Ramos B cells in control mice, whereas treatment with either 9G4 CAR-T cells or 9G4 cTCR-T cells controlled target B-cell outgrowth **(Fig. 6C-D)**. Bioluminescence from B cells was significantly reduced in both anti-9G4 T cell treatment groups compared with control T cells **(*P*<0.0001)**, with no difference between 9G4 CAR-T cells and 9G4 cTCR-T cells **(*P*=0.30) (Fig. 6E)**.

**Figure 6.**
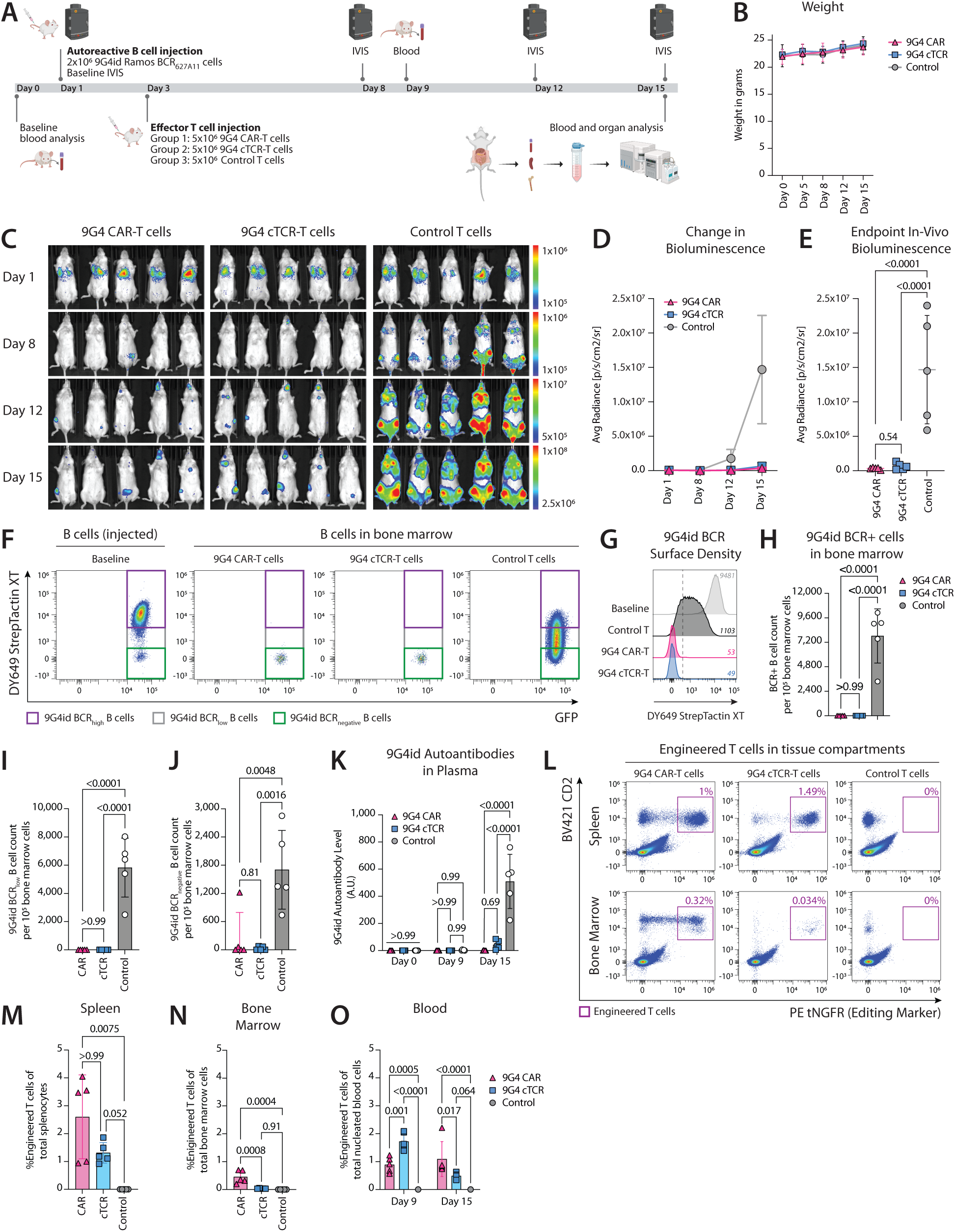
Engineered anti-9G4 T cells deplete autoreactive 9G4id Ramos B cells in vivo. **(A)** Experimental summary. NSG mice were injected with 2×10^6^ luciferase+ GFP+ SLE 9G4id Ramos BCR_627A11_ B cells, followed by treatment with 5×10^6^ 9G4 CAR-T cells, 9G4 cTCR2-T cells, or control (TCR KO) T cells; time points for IVIS imaging, blood collection, and endpoint blood and organ analyses are indicated. **(B)** Weight of mice over the experimental period. **(C)** Bioluminescence imaging (BLI) at the indicated timepoints for each treatment group (9G4 CAR-T cells [left panel], 9G4 cTCR2-T cells [middle panel], and control T cells [right panel]). **(D)** Change in BLI signal (average radiance) over time for all experimental groups. **(E)** Quantification of B cell burden by in vivo bioluminescence at experimental endpoint. **(F)** Representative flow plots of B cell frequencies and BCR surface densities at the time of injection and at the end of the experiment; gates show autoreactive 9G4id BCR_high_, 9G4id BCR_low_, 9G4id BCR_negative_ B cells, as detected by surface staining of Strep-tag II. **(G)** Histogram showing 9G4id BCR surface densities on B cells at the time of injection (baseline) and at the end of experiment in bone marrow. The median fluorescence intensity (MFI) of StrepTactin XT is given for each condition. **(H)** Quantification of live 9G4id BCR+ B cell counts (sum of BCR_high_ and BCR_low_ B cell counts) per 10^5^ bone marrow cells by flow cytometry. **(I-J)** Quantification of live 9G4id BCR_low_ B cell counts (I) and live BCR_negative_ (i.e., undetectable 9G4id BCR expression) B cell counts (J) per 10^5^ bone marrow cells by flow cytometry. **(K)** 9G4id autoantibody levels in mouse plasma at baseline and at indicated timepoints as quantified by ELISA. **(L)** Exemplary flow plots showing engineered T cells (CD2+tNGFR+) in the spleen and bone marrow at experimental endpoint. **(M–N)** Quantification of live engineered T cells as a percentage of total splenocytes (M) or total bone marrow cells (N). **(O)** Quantification of engineered T cells in blood (percent engineered T cells of total nucleated blood cells) at days 9 and 15. Each symbol represents an individual mouse; Data are shown as mean ± SD. One-way ANOVA with Tukey’s multiple comparisons were utilized in E, G, K. One-way ANOVA with Kruskal-Wallis’ multiple comparisons were utilized in J. Two-way ANOVA with Šídák’s multiple comparisons were utilized in H and L.

To validate these findings on the tissue level, we quantified autoreactive B cells in peripheral blood, the spleen, and the bone marrow by flow cytometry. Mice treated with either 9G4 CAR-T cells or 9G4 cTCR-T cells showed marked depletion of GFP+ B cells in spleen, bone marrow, and blood as compared to control T cells **(*P*<0.005; Fig. S6A-D)**, but a small percentage of GFP+ B cells was detectable in the bone marrow (0.26% of total bone marrow cells [range, 0.00-1.25%] for CAR-T cell group vs 0.03% [range, 0.00-0.084%] for cTCR-T cell group). We next specifically quantified 9G4id BCR+ B cells by interrogating the expression of Strep-tag II (part of the engineered autoreactive BCR) on the B cell surface **(Fig. 6F-G; Fig. S6H)**. Compared to control mice, both 9G4 CAR-T cells and 9G4 cTCR-T cells potently depleted 9G4id BCR+ B cells in bone marrow **(*P*<0.0001; Fig. 6H)**, spleen **(*P*=0.0026; Fig. S6I)**, and peripheral blood **(*P*<0.0001; Fig. S6J)**, with comparable potency between the two experimental anti-9G4 T cell groups **(*P*>0.99; Fig. 6H, S6I–J)**. Indeed, residual B cell signals observed at the end of the experiment were due to growth of BCR-negative B cells that represented ∼1% of injected Ramos B cells. Notably, Ramos B cells showed a propensity to markedly downregulate their surface BCR density in vivo even in the absence of selective pressure **(Fig. 6F-G, control)**. Interestingly, these B cells with low, or by flow cytometry even undetectable, surface expression of autoreactive BCRs were still effectively killed by anti-9G4 T cells **(P<0.005; Fig. 6I-J)**. Consistent with their specific cytotoxicity in vivo, plasma levels of 9G4id autoantibody increased in control mice but were effectively suppressed following treatment with anti-9G4 T cells (*P*<0.0001; 9G4 CAR-T cells vs 9G4 cTCR-T cells, *P*=0.69) **(Fig. 6K).**

We next assessed the biodistribution and persistence of engineered T cells in vivo. CD2+ T cells, chosen as a marker for T cells as CD3 surface expression is lost in 9G4 CAR-T cells after knock-out of *TRAC*, were detectable in all groups over the course of the experiment **(Fig. S6A)**. Across tissues, mice receiving 9G4 CAR-T cells showed higher frequencies of CD2+ T cells compared with the 9G4 cTCR-T cell and control T cell groups **(Fig. S6E-G)**. Engineered anti-9G4 T cells, identified by co-expression of CD2 and tNGFR, showed persistence in mice and remained detectable in peripheral blood, spleen, and bone marrow throughout the experiment **(Fig. 6L, S6K)**. The frequencies of engineered T cells varied across treatment groups. While 9G4 CAR-T cells showed higher abundance the in the bone marrow **(*P*=0.0008; Fig. 6N)**, the blood **(*P*=0.017; Fig. 6O)**, and a trend toward higher abundance the in the spleen **(*P*>0.99; Fig. 6M)** at day 15, 9G4 cTCR-T cells showed more pronounced expansion in the peripheral blood at day 9 **(*P*=0.001) (Fig. 6O, S6K)**, indicating different temporal dynamics of expansion. Collectively, these data show that both 9G4 CAR-T cells and 9G4 cTCR-T cells effectively killed 9G4id Ramos B cells in vivo and reduced circulating 9G4id autoantibody.

## Discussion

Cellular therapies that indiscriminately target B cells can achieve drug-free disease remission in patients with severe autoimmune diseases, but the risk of serious adverse events, including infection and cytokine-related toxicities (*63*), limits their widespread use. Precision cellular therapies engineered to selectively target pathogenic B cell subsets are therefore being explored to harness the potency of immune effector cell therapies without compromising protective immunity (*24*). If successful, such approaches could move immune effector cells from salvage care to safer, frontline interventions that transform treatment across the spectrum of autoimmune disease.

One such approach comprises antigen-specific cellular immunotherapies that are engineered to target autoreactive B cells directly through binding of their cognate autoantigen. These strategies, however, are only suitable for the treatment of autoimmune diseases in which one autoantigen is the primary target of the pathogenic B cell response. In SLE, the broad autoreactome targeted by pathogenic B cells makes antigen-specific approaches unfeasible (*25*). To overcome this, we developed a precision cellular therapy approach for the treatment of SLE that instead targets the 9G4id present in *VH4-34*-derived BCRs, exploiting a shared origin and structural feature of autoreactive B cells in SLE (*35, 36*), including those cross-reactive with dsDNA, DNAse1L3, Sm/RNP, Ro52, and cardiolipin (*28, 35, 36*). To this end, we evaluated the efficacy and specificity of T cells expressing different synthetic immune receptor formats, including CARs and cTCRs, for the targeting autoreactive 9G4id B cells. We found that all engineered anti-9G4 T cells showed high specificity and eliminated 9G4id B cells without depleting non-9G4 B cells in vitro. This was observed regardless of somatic hypermutation, reactivity, or disease origin (i.e., SLE, CAD, or lymphoma) of the targeted B cells. Notably, 9G4id B cells, whether engineered B cell lines or primary B cells from patients with SLE, were efficiently eliminated despite their capacity to secrete autoantibodies, including under conditions of exogenous stimulation of antibody secretion. These findings indicate that soluble autoantibodies did not impair targeting of BCR-specific B cells across different therapeutic strategies, as demonstrated in pemphigus vulgaris (*20*). Consistent with these findings, both 9G4 CAR-T cells and 9G4 cTCR-T cells effectively eliminated autoreactive 9G4id Ramos B cells in vivo and reduced circulating 9G4id autoantibody levels, including when targeting B cells with low surface BCR density. The deep, selective depletion of autoreactive B cell subsets, as compared to the broad targeting of B cells, represents a major step towards immune effector cell therapies that do not increase the risk of infection.

Adapting these synthetic immune receptors in ways that obviate conditioning therapy, which by itself causes transient immunosuppression (*64*), is a natural evolution of this approach. While the reduction or elimination of conditioning therapy in the treatment of autoimmune diseases is currently being investigated for ex-vivo engineered cellular therapies (*65*), in-vivo delivery of mRNA or DNA encoding chimeric immune receptors to T cells represents an emerging opportunity to advance precision cellular therapies without even transient immunosuppression (*66, 67*). These approaches allow for the direct generation of engineered T cells within the patient, thereby eliminating the need for complex ex-vivo manipulation and obviating the need for conditioning therapy. In patients with SLE, transient expression of synthetic immune receptors in T cells may be sufficient to ameliorate or control disease without risk of long-term immune system alterations or secondary malignancy (*68, 69*). Leveraging in-vivo delivery of mRNA to transiently introduce anti-9G4 synthetic immune receptors may ultimately present a safe, scalable, and clinically sufficient option for patients with mild to moderate SLE, in whom the risks associated with conditioning therapy (e.g., infection, infertility), broad B cell depletion (infection), and currently unquantifiable long-term risks of gene editing (e.g., secondary cancers) pose major barriers to treatment with existing cellular therapies (*69–71*).

Cytokine-related toxicities, including CRS, ICANS, and delayed cytopenia (*18*), are another major barrier to administering T cell therapies at scale, necessitating close monitoring and early therapeutic intervention. While these toxicities appear to be less severe in patients with autoimmune diseases than in patients with B cell cancers (*72*), CRS remains common (*14, 15, 73, 74*) and high-grade neurotoxicity has been reported (*75, 76*). Minimizing cytokine release is therefore essential to improve the safety of cellular therapies, enable their use in outpatient settings, and extend their application to patients with non-life-threatening autoimmune disease. One opportunity to mitigate the risk of cytokine-related toxicities is to substantially reduce the target cell burden, a benefit common to all precision cellular therapies that target only a small fraction of total B cells (*77, 78*). This theoretical benefit was observed in our study where direct comparison of matched-donor anti-9G4 T cells and CD19 CAR-T cells against SLE PBMCs showed significantly reduced cytokine release in the former case. Cytokine release may also be curtailed by strategic selection of effector cell types and phenotypes, as well as through optimization of immune receptor architecture and signaling components (*79*). In studying different synthetic immune receptor formats in direct comparison, we found that 9G4 CAR-T cells demonstrated the highest release of IFN-γ, while cytokine release was markedly reduced when using cTCR-T cell formats. These differences may be explained by the co-stimulatory domain (i.e., CD28) in the CAR constructs, in addition to fundamental differences in receptor regulation and immune synapse formation between CARs and TCRs (*42, 47, 80–85*). 9G4 CAR-T cells exhibited antigen-independent expansion, consistent with constitutive signaling, which was importantly not observed in 9G4 cTCR-T cells. The differences in cytokine release between the immune receptor scaffolds were more pronounced when targeting 9G4id Ramos B cells, derived from transformed B cells of a patient with Burkitt lymphoma, than in co-cultures with SLE patient PBMCs. This may reflect several factors, including differences in target cells (e.g., resistance to cytotoxic killing pathways in transformed vs. primary B cells), differences in the functional state of effector T cells in SLE patients vs. healthy donors (*86*), or a combination thereof. While co-stimulatory domains are essential for durable responses in cancer, the need for long-term T cell persistence for durable remission in autoimmune disease is unproven, and persistence beyond deep B cell depletion (conceptually referred to as “immune reset” in the current literature) may in fact be detrimental. We observed that 9G4 cTCR-T cells expanded earlier in mice and were less abundant in the spleen, bone marrow, and peripheral blood than 9G4 CAR-T cells at the end of treatment. The reduced T cell proliferation and cytokine secretion associated with 9G4 cTCR-T cells did not compromise in-vivo efficacy in our model. Indeed, 9G4 CAR-T cells and 9G4 cTCR-T cells eliminated 9G4id B cells and circulating autoantibodies with comparable potency in vivo. The combination of preserved efficacy and reduced cytokine release suggests that 9G4 cTCR-T cells may offer a favorable safety profile for use in patients with mild-to-moderate SLE or other autoimmune diseases, in whom engineered T cell persistence is not a goal.

Disease heterogeneity is a significant barrier to therapeutic interventions in SLE (*87*). While 9G4id B cells are a major source of pathogenic autoantibodies in lupus (*28, 35, 36*), it is unknown whether their deep depletion alone would suffice to ameliorate disease in all patients. The contribution of 9G4id B cells to organ damage likely varies among individuals, highlighting the need for diagnostics to identify patient subsets most likely to respond to precision immunotherapies. Such diagnostics may include BCR repertoire sequencing (to quantify VH4-34 clonal expansion) (*88, 89*), flow cytometry to enumerate class-switched 9G4id memory B cells (*33*), or immunoassays measuring levels of 9G4id autoantibodies longitudinally (*28*). Notably, autoreactive B cell clones outside the VH4-34 repertoire may contribute to disease. For instance, 9G4id antibody depletion in serum reduced anti-DNASE1L3 autoantibodies by up to 80% in SLE patients (*28*). Thus, it is conceivable that precision therapies targeting 9G4id B cells may control disease in some patients, while others may only achieve partial responses. Future research focused on comprehensively defining the contribution of 9G4id B cells in heterogenous disease cohorts of SLE will be critical to advance precision medicines for lupus.

The utility of 9G4-targeted therapies extends beyond SLE and could be of value in other B cell-driven diseases. One notable condition is cold agglutinin disease (CAD), a type of autoimmune hemolytic anemia where IgM autoantibodies bind to red blood cells at low temperatures, leading to their destruction (*90*). These IgM autoantibodies are almost invariably encoded by the VH4-34 gene, making 9G4id B cells a promising target for therapeutic intervention (*55, 56*). We and others have identified clonally expanded 9G4id B cells in the CSF of patients with multiple sclerosis (MS) (*89*), and similar contributions to autoantibody compartments are found in other autoimmune diseases (*91*), an opportunity for their precision targeting in select patients guided by molecular diagnostics. Finally, 9G4-targeted immunotherapies may be promising precision therapeutics in patients with various B cells cancers which are abnormally enriched in VH4-34 usage (*59*). We expect that the selection and utility of different synthetic immune receptor designs will be dictated by the patient population, and benefits observed for the treatment of cancer may not be equally applicable for patients with autoimmune diseases.

## Materials and Methods

### Study Design

The goal of the study was to develop and evaluate synthetic immune receptor-T cell therapies for the precision targeting of 9G4id B cells in patients with SLE, aiming to eliminate pathogenic 9G4id B cells while sparing most normal B cell populations. Specifically, this study comparatively tested anti-9G4 CAR and anti-9G4 cTCR designs for the treatment of autoimmune diseases. Primary human T cells from healthy donors or patients with SLE were genetically modified using CRISPR-Cas12a to eliminate endogenous TCR expression and introduce anti-9G4 CAR or anti-9G4 cTCR constructs via homology-directed repair (HDR). Synthetic immune receptor expression in T cells was quantified by flow cytometry, and successfully edited T cells were enriched by positive selection. Ramos B cell lines were engineered using CRISPR-Cas9 HDR to replace endogenous BCR expression with patient-derived monoclonal BCRs from patients with SLE, cold agglutinin disease (CAD), or irrelevant BCRs, creating isogenic cell lines with and without 9G4id expression for specificity testing. T cell-mediated cytotoxicity was measured using live-cell imaging, flow cytometric enumeration of target cells, and (auto)antibody production. T cell activation was assessed by quantification of cytokine and chemokine secretion using enzyme-linked immunosorbent assay (ELISA) and Meso Scale U-PLEX. T-cell proliferation was interrogated through fluorescent dye dilution by flow cytometry. The function of engineered, donor-matched T cells to deplete primary 9G4id B cells from SLE patients was tested in PBMCs using both B cell FluoroSpot and bulk BCR repertoire sequencing. Comparative analyses between CAR-T and cTCR-T cell platforms were performed to evaluate potency, cytokine release, antigen-dependent and antigen-independent proliferation. Statistical tests, number of replicates, and number of experiments are listed in the figure legends.

### Synthetic immune receptor design and generation

Nucleotide sequences encoding 9G4 cTCR, 9G4 CAR, and CD19 CAR constructs were designed in silico. Sequences were codon optimized for human expression, de-novo synthesized (GeneArt), and cloned into pUC19-derived vectors containing an EF1α promoter, Kozak sequence, signal peptide, synthetic immune receptor domain(s), 2A ribosomal skip sequence, signal peptide, truncated nerve growth factor receptor (tNGFR), stop codon, and polyadenylate terminator, all flanked 5’ and 3’ by homology arms (HA) for the human *TRAC* locus. For cTCRs, fully human TCRα and TCRβ chains separated by 2A sequences were linked to either anti-9G4 scFv via an EAAAK linker to TCRα (cTCR1), or by fusing VH and VL fragments of the anti-9G4 antibody to the TCRα and TCRβ chains (cTCR2), respectively. CARs utilized second-generation CAR designs comprising a CD8α signal peptide, various hinge domains, transmembrane domain, a CD28 co-stimulatory domain, and a CD3ζ signaling domain. Homology-directed repair templates (HDRTs) for CRISPR editing were generated by PCR amplification from vector templates (Q5 Hot Start High-Fidelity 2X Master Mix, New England BioLabs), purified using AMPure XP Reagent (Beckman Coulter, A63880), eluted in nuclease-free water, and quantified using a NanoDrop spectrophotometer (Thermo Fisher).

### Gene editing of primary human T cells

Peripheral blood mononuclear cells (PBMCs) were isolated from healthy donor leukopaks (STEMCELL) by Ficoll-Paque PLUS (Cytiva) gradient centrifugation and cryopreserved in CryoStor CS10 (BioLife Solutions). For some experiments, PBMCs were collected from patients with SLE under a Johns Hopkins Medicine IRB approved protocol (IRB00307779), cryopreserved in CryoStor CS10, and thawed for isolation and editing. CD3+ T cells were isolated from PBMCs by negative immunoselection (STEMCELL, 17951) and resuspended in RPMI-1640 (ATCC, 30-2001) supplemented with 10% FBS (Gibco, A5669701), 1% penicillin-streptomycin (P/S, Thermo Fisher, 15140163), 100 IU/mL recombinant human interleukin (IL)-2 (Proleukin, Prometheus Laboratories), and 5 ng/mL recombinant human IL-7 (BioLegend, 581908), hereafter called T-cell media. T cells were immediately activated with Dynabeads Human T-Activator CD3/CD28 (Thermo Fisher, 11132D) at a 1:1 bead-to-cell ratio for 48 hours, after which beads were removed with a magnet. To edit T cells, 50 pmols of Cas12a sgRNAs targeting either *TRAC* or *TRBC* were incubated with 25 pmols of Alt-R Cas12a (Cpf1) Ultra (IDT, 10001273) and Alt-R Cpf1 Electroporation Enhancer (IDT, 1076301) in Nuclease Free Duplex Buffer (IDT, 11-01-03-01) for 15 minutes to form RNPs, before combining both RNPs at a 1:1 volume ratio. *TRAC*- and *TRBC*-targeted RNPs were then incubated with 0.5 µg of cTCR/CAR HDRTs diluted in OptiMEM (Gibco). For each condition, 1×10^6^ T cells were pelleted at 90 x *g* for 10 minutes, resuspended in 20 µL P3 buffer (Lonza, V4XP-3032), and combined with 5 µL of Cpf1 RNP/HDRT complex. T cells were nucleofected in 16-well cuvettes (Lonza, V4XP-3032) using a 4D Nucleofector X-Unit (Lonza, AAF-1003X) and pulse code EH115. After nucleofection, 80 µL pre-warmed media without cytokines (RPMI-1640, 10% FBS, 1% P/S) was added to cells, and the cuvette strip was placed at 37°C, 5% CO_2_ for 30 min. The recovered cells were then resuspended in T-cell media and incubated in 24-well plates at 37°C, 5% CO_2_. Media was changed every 2-3 days, and T cells were expanded for at least 11 days until functional assays were performed. All sgRNA sequences are listed in the data supplement.

### Generation of autoreactive human B cell lines

Ramos RA1 B cells (CRL-1596, ATCC) were cultured in RPMI-1640 (ATCC) supplemented with 10% FBS and 1% P/S, hereafter called Ramos media, in a humidified incubator at 37°C and 5% CO_2_. Cells were transduced with IVISbrite Red F-luc-GFP (BD) lentiviral particles to enable the stable expression of green fluorescent protein (GFP) and firefly luciferase (Luc), and edited cells isolated by FACS using a BD FACSMelody Cell Sorter. To generate autoreactive 9G4id B cell lines, Ramos RA1 cells (“wild-type”) were engineered using CRISPR/Cas9 homology-directed repair to express desired monoclonal BCRs and antibodies. Coding sequences of SLE patient-derived, autoreactive 9G4id BCRs (clones 627A11, 75G12, 88F7)(*28*), CAD patient-derived, autoreactive 9G4id BCRs (cold-agglutinin antibody clones KAU(*92*), FS-1 and FS-2(*93, 94*)), and non-9G4 BCRs using VH4-4 were introduced into the *IGH* locus to replace the Ramos cell’s endogenous BCR and to generate autoreactive Ramos B cells that express both soluble and membrane-bound immunoglobulin under control of a locus-specific promoter, while preserving natural splicing(*49*). Briefly, nucleotide sequences comprising IgVH promoter, signal peptide, immunoglobulin variable light (VL), constant light (CL), Strep-tag II linker, and variable heavy (VH) chains, flanked by homology arms for the human *IGH* locus, were de-novo synthesized (GeneArt) and cloned into suitable vectors. HDRTs were generated as described above. To edit Ramos cells, 50 pmol of purified Cas9 nuclease (Alt-R SpCas9 Nuclease V3, IDT) was combined with 100 pmol of *IGH*-targeted Cas9 sgRNA (IDT, custom) and Alt-R Cas9 Electroporation Enhancer (IDT, 1075915) for 15 minutes at room temperature (RT). RNPs were then incubated with 60 pmol of HDRTs diluted in OptiMEM. For each condition, 1×10^6^ Ramos cells were resuspended in 20 µL SF buffer (Lonza, PBC2-00675) and combined with 5 µL of Cas9 RNP/HDRT complex. Ramos cells were nucleofected in 16-well cuvettes using a 4D Nucleofector X-Unit and pulse code CV-104. After nucleofection, 80 µL pre-warmed Ramos media was added to cells, and the cuvette strip was placed at 37°C, 5% CO_2_ for 30 min. The recovered cells were then resuspended in Ramos media and incubated in 24-well plates at 37°C, 5% CO_2_. After one week, edited cells were enriched by one or more rounds of positive selection using StrepTactin magnetic beads (iba, 2-5090-010). Edited cells were isolated by FACS based on robust GFP and engineered BCR expression, stained with DY649-StrepTactin XT (iba, 2-1568-050). Sorted cells were plated by limiting dilution to obtain individual autoreactive B cell clones. All sgRNA sequences are listed in the data supplement.

### Flow cytometry

Flow cytometry was performed using an Attune NxT cytometer (Thermo Fisher). Cells of interest were labeled with a viability dye, either LIVE/DEAD Fixable Near-IR Dead Cell Stain (Thermo Fisher, L34975) or LIVE/DEAD Fixable Violet Dead Cell Stain (Thermo Fisher, L34955). T cells were stained with combinations of the following anti-human antibodies: APC anti-CD3 (clone SK7, BioLegend, 344812), Brilliant Violet (BV) 421 anti-CD3 (clone SK7, BioLegend, 344834), FITC anti-CD3 (clone SK7, BioLegend, 344804), APC anti-human CD271 (clone ME20.4, BioLegend, 345108), FITC anti-human CD271 (clone ME20.4, BioLegend, 345104), FITC anti-PD-1 (clone NAT105, BioLegend, 367412), APC anti-PD-1 (clone EH12.2H7, BioLegend, 329908), APC anti-CD8 (clone SK1, BioLegend, 344722), BV421 anti-CD8 (clone RPA-T8, BioLegend, 301036). The expression of cTCRs and CARs was assessed using DyLight 488-conjugated (Abcam, ab201799) 9G4id IgG1 (clones 627A11, 75G12, 88F7), expressed in Expi293 cells (Gibco) and purified as described(*28*). When staining human PMBCs, Human TruStain FcX (BioLegend, 422302) was used. Ramos B cells were stained with combinations of the following anti-human antibodies: Alexa Fluor (AF) 488 anti-CD20 (clone 2H7, BioLegend, 302316), DyLight 650 anti-9G4 (conjugated in house; Abcam, ab201803), DY-649 Strep-Tactin XT (iba, 2-1568-050). Ramos cells were stained with AF488 anti-CD19 (clone SJ25C1, BioLegend, 363038), BV421 anti-CD19 (clone SJ25C1, BioLegend, 363018), AF488 anti-CD20 (clone 2H7, BioLegend, 302316), BV421 anti-CD20 (clone 2H7, BioLegend, 302330), idiotypic rat anti-human 9G4 (DyLight 650 conjugated). Flow cytometry data were analyzed with FlowJo v.10.10.1 software (BD).

### Mouse xenograft models

Female NOD.Cg-Prkdcscid Il2rgtm1Wjl/SzJ (NSG) mice (6-8 weeks of age) were obtained from The Jackson Laboratory (JAX, 005557) or the Johns Hopkins Sidney Kimmel Comprehensive Cancer Center Animal Resources facility and housed in accordance with an Institutional Animal Care and Use Committee-approved protocol at The Johns Hopkins University. 9G4id Ramos BCR_627A11_ B cells and human T cells were administered via tail-vein injection in 200 μL RPMI 1640; injection schedules are shown in Fig. 6. B-cell growth was monitored by bioluminescence imaging, and body weights were recorded at indicated time points. For imaging, mice received an intraperitoneal injection of 150 μL RediJect D-Luciferin Ultra (PerkinElmer, 770505), were anesthetized with inhaled isoflurane, and were imaged using an IVIS system (PerkinElmer). Peripheral blood was collected in EDTA microvettes (Sarstedt, NC9299309) from the submandibular vein. At endpoint, spleen and bone marrow cells were collected.

### Processing of blood, spleen, and bone marrow for flow cytometric analysis

EDTA whole blood (100 μL) was subjected to red blood cell lysis using ACK buffer (Quality Biological, 118-156-721) and washed with PBS. Spleens were harvested into ice-cold RPMI 1640 and mechanically dissociated by gently pressing tissue through a 40-μm cell strainer (Alkali Scientific, TS40) using a syringe plunger. Cells were washed, pelleted, and subjected to ACK-mediated red blood cell lysis; lysis was quenched with DPBS containing 2% BSA. For bone marrow cells, femurs were excised, cleaned of surrounding tissue, and cut at both ends. Marrow was flushed with ice-cold PBS through a 40-μm cell strainer, and any clumps were gently disrupted using a syringe plunger. Cells were pelleted, subjected to ACK red blood cell lysis, quenched as above, washed, and resuspended for flow cytometric staining.

### In-vitro co-culture of engineered B cell lines

Anti-9G4 cTCR-T cells, CAR-T cells, or control T cells (“effector cells”) were combined with 1×10^4^ autoreactive or unedited GFP+ Luc+ Ramos B cells (“target cells”) at defined E:T ratios in T-cell media, unless specified otherwise. Co-cultures were incubated for 2.5-4.5 days at 37°C, 5% CO_2_, humidified air in tissue culture-treated 96-well microplates (Costar, Corning), as specified in the figure legends. Longitudinal killing of GFP+ Ramos B cells was quantified every 4 hours using an Incucyte SX5, G/O/NIR module, and 10-20x objective (Sartorius). Green integrated intensity for each well normalized to day 0 was used for statistical analysis. Endpoint flow cytometry analysis of samples was conducted by gating on live, single GFP+ B cells. Viability was calculated by normalizing to the absolute B cell number in the control T cell groups using the formula number_experimental well_/ number_control well_ x 100. At assay endpoint, conditioned co-culture media was also collected and stored for cytokine assays.

### Repeated stimulation assays

To evaluate response to multiple stimulations of autoreactive target cells, engineered T cells (5×10^4^) were combined with GFP+ Ramos B cells (1×10^4^) at a 5:1 E:T ratio in 200 µL of T-cell media, and cells were imaged on a live-cell imager (Incucyte) as described above. At 3 days, 100 µL of conditioned supernatant was removed, and additional Ramos B cells (2×10^4^ cells in 100 µL of new media) were added to the co-culture. At 6 days, additional Ramos B cells (4×10^4^ cells) were added as described previously. At the assay endpoint (12 days), live, single Ramos B cells and engineered T cells were quantified by flow cytometry. T cells proliferation was assessed by flow cytometry using Cell Trace Violet (ThermoFisher, C34557). The weighted average of generations was calculated using AVG = (N_0_*0 +… + N_g_*g)/N_total_, where g represents each generation number, and N represents the total cell events within that generation’s gate.

### In-vitro co-culture of human PBMCs

Cryopreserved PBMCs (1×10^5^ viable cells) collected from patients with SLE were thawed and combined with engineered, donor-matched human T cells (5×10^3^ viable cells) in serum-free media (ImmunoCult-XF B Cell Base Medium, supplemented with ImmunoCult-ACF Human B Cell Expansion Supplement, STEMCELL) or RPMI-1640 media (ATCC) supplemented with 10% FBS, 1% P/S, and human IL-2 and IL-7, as specified in figure legends. After 48 hours, conditioned co-culture media was collected and stored for cytokine assays, and new serum-free media was added. After another 48-96 hours of B cell activation, cells were collected and subjected to B cell FluoroSpot assay (Mabtech) and mRNA-based IgH repertoire sequencing (iR-RepSeq+, iRepertoire).

### Autoantibody ELISAs

9G4id antibody levels were quantified in cell culture supernatants and mouse plasma using an in-house ELISA. Briefly, polystyrene plates (Costar, 9018) were coated overnight at 4°C with purified rat anti-9G4 antibody (IGM Biosciences) in PBS (pH 7.4) at 100 ng/well for cell culture supernatants or at 200 ng/well for mouse plasma; wells coated with PBS alone served as background controls. Coated plates were washed with PBS 0.05% Tween-20 (PBS-T, Thermo Fisher) and remaining binding sites blocked with PBS-T containing 5% non-fat dry milk (PBS-TM) for 1 hour at RT. Culture supernatants (1:10) and mouse plasma (1:20) were diluted in PBS-TM 1%, and 100 <u>μL</u> was added to washed plates, and incubated for 1 hour at RT, 150 rpm. Binding of human 9G4id antibodies was detected using HRP-conjugated goat anti-human IgM (Jackson ImmunoResearch, 109-035-129), diluted 1:2,000 in PBS-TM 1% for cell culture supernatants or diluted 1:10,000 in PBS-TM 1% for mouse plasma, incubated for 1 hour at RT, 150 rpm, protected from light. TMB peroxidase substrate solution (SureBlue, KPL) was added to wells, and the reaction was stopped with 1M H_2_SO_4_ (Thermo Fisher). Absorbance was measured at 450 nm (background 620 nm) using an Absorbance 96 microplate reader (Byonoy). Antibody levels were calculated in reference to wells coated with a serial dilution of native human IgM (Abcam, ab91117) that served as a standard. Anti-dsDNA antibodies in cell culture supernatants (1:5 diluted) were measured using a modification of the QUANTA Lite dsDNA ELISA (Inova, 708510), replacing the secondary antibody with HRP-conjugated goat anti-human IgM at 1:10,000 dilution.

### FluoroSpot assay

9G4id antibody-secreting B cells in PBMCs from patients with SLE were measured using a modified B cell FluoroSpot assay. FluoroSpot plates (MabTech, X-05R-10) were coated with 1500 ng/well of rat anti-9G4 antibody or 1500 ng/well anti-human IgG (MT91/145, MabTech, 3850-3) in PBS pH 7.4 (Gibco), overnight at 4°C. Plates were washed with PBS, and the membrane blocked with PBS containing 5% bovine serum albumin (BSA) for 1 hour at RT. PBMCs treated with engineered T cells or control T cells were added at 5×10^4^ or 5×10^3^ live cells/well and incubated for 16-20 hours at 37°C, 5% CO_2_. Plates were washed to remove cells and membranes incubated with anti-human IgG 550 (MT78/145, MabTech, 3850-5R) in PBS 0.1% BSA for 2 hours at RT. Fluorescence enhancement solution (MabTech, 3641-F10-60) was then added to wells. After final washes in PBS pH 7.4 (Gibco), membranes were air dried and stored in the dark. Fluorescent spots corresponding to total IgG and 9G4id IgG ASCs were quantified using an ImmunoSpot Analyzers (CTL ImmunoSpot S6 Universal M2 Analyzer, S6UNV2) at excitation 550 nm/ emission 600 nm.

### Cytokine assays

Cytokines were quantified in cell culture supernatants using an IFN-γ ELISA (R&D Systems) or U-PLEX 10-plex cytokine assay for human IL-2, IL-4, IL-6, IL-8, IFN-γ, IL2Rα, granzyme A, granzyme B, TNF-α, TNF-β, GM-CSF (Meso Scale Diagnostics, MSD, K151AEL-1). Samples, standards, or controls were added at 50 µL/well (MSD) or 100 µL/well (Human IFN-γ ELISA) and plates incubated for 2 hours at RT, 120 rpm. Plates were washed three times using PBS pH=7.5 0.05% Tween 20. Pre-diluted secondary antibody was added at 50 µL/well (MSD) or 100 µL/well (Human IFN-γ ELISA), and plates incubated for 1 hour at RT, 120 rpm. Following final washes, MSD Read Buffer was added at 150 µL/well and plates read using a Sector Imager 2400 (MSD). Raw electrochemiluminescence data were analyzed using the Discovery Workbench 3.0 software (MSD). A 4-parameter logistic (4PL) curve fit was generated for each analyte using standard data points to determine the concentrations of unknown samples.

### BCR repertoire sequencing and analysis

For BCR repertoire analysis, cells from co-culture experiments were washed in PBS, pelleted, and stored at −80°C in Buffer RLT (Qiagen, 79216). mRNA was extracted using RNeasy RNA Extraction kits (Qiagen). Samples were amplified using BCR heavy chain (IGH) primers, and adaptome amplification was performed using iR-RepSeq+ (iRepertoire, Inc.). Next-generation sequencing libraries were generated for each sample. Unique molecular identifiers (UMIs) were incorporated during the reverse transcription step to distinguish individual RNA molecules. Reverse transcription was performed using OneStep RT-PCR mix (Qiagen) with C-gene primer mix, followed by selection of first-strand cDNA selection and removal of remnant primers via SPRIselect bead purification (Beckman Coulter). A second round of amplification was conducted using a V-gene primer mix, followed by SPRIselect bead purification. Library amplification was performed with primers targeting communal sites engineered onto the 5′ ends of the C- and V-primers. The final libraries contain Illumina dual-index sequencing adapters, a 10-nucleotide random region, and an 8-nucleotide internal barcode associated with the C-gene primer. Sequencing coverage includes from within the framework 1 region to the C-region, including CDR1, CDR2, and CDR3. BCR amplified libraries were multiplexed, pooled, and sequenced on 20% of a Nextseq 1000 P1 600 cycle. RepSeq+ sequencing raw data were analyzed using iRmap. Briefly, sequence reads were de-multiplexed according to Illumina dual indices and barcode sequences. Merged reads were mapped to germline V, D, J, and C reference sequences using an IMGT reference library. CDR3 regions were identified, extracted, and translated into amino acids. The dataset was condensed by collapsing UMIs and CDR3 sequences to correct for sequencing and amplification errors. Reads sharing identical CDR3 and UMI combinations were condensed into a single UMI count. Repertoire analysis including diversity metrics, V(D)J usage, and clonal expansion was performed using Immunarch R package. The chord diagram was graphed using Circlize R(*95*) package.

### Statistical analyses

Statistical tests were applied to experimental group data as outlined in the figure legends and dataset. In general, comparisons were made between the control conditions (e.g., target B cells not expressing 9G4id) vs. the experimental conditions (e.g., target B cells expressing 9G4id BCRs). *P* values <0.05 were considered significant, unless otherwise specified. Analyses were performed using GraphPad Prism v10.

## Acknowledgments

We thank Armaan Bahl (Johns Hopkins), Ali Dbouk (Johns Hopkins), Jacqueline Douglass (Johns Hopkins), Jiaxin Ge (Johns Hopkins), Alisha Mills (Johns Hopkins), Tushar Nichakawade (Johns Hopkins), Nickolas Papadopoulos (Johns Hopkins), Drew Pardoll (Johns Hopkins), and Evangeline Watson (Johns Hopkins) for their scientific and technical support. We thank Christopher Thoburn for technical support, the SKCCC Immune Monitoring Core (Johns Hopkins), and the Bloomberg Flow Cytometry and Immunology Core. We thank Freda Stevenson for her expertise and scientific support. We thank our patients for contributing blood samples to the biorepository at Johns Hopkins.

## Author Contributions

J.L. and M.F.K. conceived and designed the research studies; J.L., Y.X., B.J.M., C.G., E.S., D.F., A.M., K.C., B.M., T.O.A., S.G., K.J.K., T.S.A., and M.F.K. conducted experiments; R.B. and I.S. cloned SLE patient antibodies (clones 627A11, 75G12, 88F7). J.L., Y.X., and M.F.K. acquired and analyzed data; S.R.D., X.L., N.M., A.H.P., C.B., S.P., K.W.K, S.Z., F.A., and B.V. assisted with analysis and interpretation of results. J.L. and M.F.K. wrote the original draft of the manuscript; all other authors reviewed, edited, and approved the manuscript.

## List of Supplementary Materials

### Supplementary Figures

**Figure S1.** Design and characterization of different 9G4 CAR-T cells

**Figure S2.** Long-term functional assessment of 9G4 synthetic immune receptor T cells in vitro

**Figure S3.** Efficacy of 9G4 CAR-T cells and 9G4 cTCR-T cells against autoreactive 9G4id B cells in cold agglutinin disease (CAD) and B cell lymphoma

**Figure S4.** Cytokine release by 9G4 CAR-T cells and 9G4 cTCR-T cells targeting Ramos B cells

**Figure S5.** 9G4 cTCR-T cells and 9G4 CAR-T cells eliminate 9G4id B cells from patients with SLE

**Figure S6.** Flow cytometric quantification of B cells and T cells in vivo

### Supplementary Tables

**Table S1.** Patient characteristics

**Table S2.** Frequency of B cells in SLE PBMCs and autologous engineered T cell properties

**Table S3.** Nucleotide sequences of CRISPR guide RNA

**Table S4.** Amino acid sequences of 9G4id and control BCRs

### Other Supplementary Materials

**Dataset.** Details and extended statistical analyses for Figures and Supplementary Figures

**Movie S1-S6.** Live-cell imaging of anti-9G4 T cells targeting 9G4id or non-9G4 Ramos B cells

