## Supplementary figures and tables for "Precision targeting of autoreactive B cells in systemic lupus erythematosus using anti-9G4 idiotope synthetic immune receptor T cells"

**The file includes:**

Figs. S1 to S6
Tables S1 to S4

**Other Supplementary Materials for this manuscript include the following:**

Data File S1 to S9 (separate file)
Dataset (separate file)
Movies S1 to S6 (separate file)

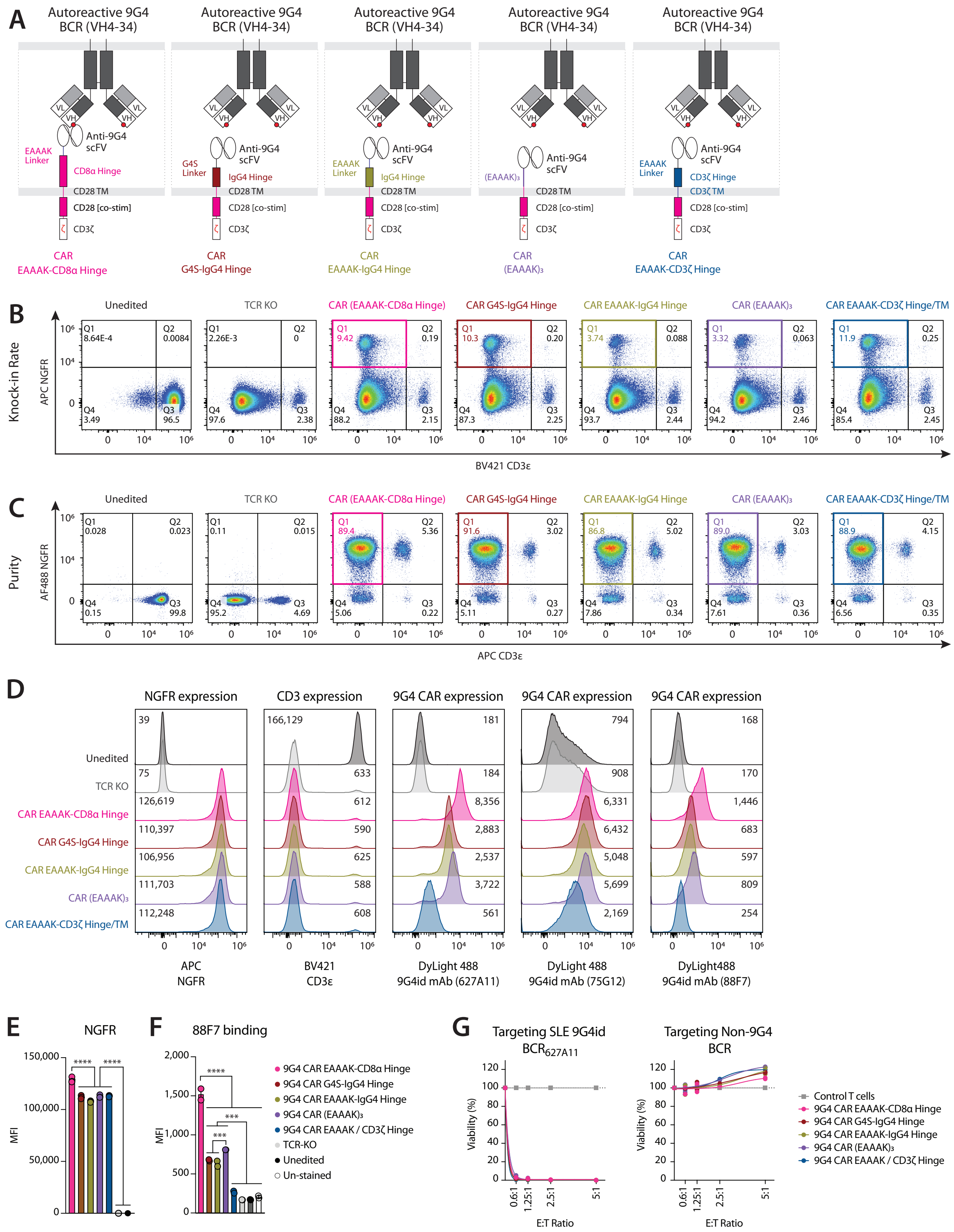
A
Autoreactive 9G4 BCR (VH4-34)
Anti-9G4 scFV
EAAAK Linker
CD8α Hinge
G4S Linker
IgG4 Hinge
(EAAAK)₃
CD3ζ Hinge
CD3ζ TM
CD28 TM
CD28 [co-stim]
CD3ζ
CAR EAAAK-CD8α Hinge
CAR G4S-IgG4 Hinge
CAR EAAAK-IgG4 Hinge
CAR (EAAAK)₃
CAR EAAAK-CD3ζ Hinge
B
Knock-in Rate
APC NGFR
Unedited
TCR KO
CAR (EAAAK-CD8α Hinge)
CAR G4S-IgG4 Hinge
CAR EAAAK-IgG4 Hinge
CAR (EAAAK)₃
CAR EAAAK-CD3ζ Hinge/TM
BV421 CD3ε
C
Purity
AF488 NGFR
APC CD3ε
D
NGFR expression
CD3 expression
9G4 CAR expression
APC NGFR
BV421 CD3ε
DyLight 488 9G4id mAb (627A11)
DyLight 488 9G4id mAb (75G12)
DyLight488 9G4id mAb (88F7)
E
NGFR
MFI
F
88F7 binding
9G4 CAR EAAAK-CD8α Hinge
9G4 CAR G4S-IgG4 Hinge
9G4 CAR EAAAK-IgG4 Hinge
9G4 CAR (EAAAK)₃
9G4 CAR EAAAK / CD3ζ Hinge
TCR-KO
Unedited
Un-stained
G
Targeting SLE 9G4id BCR627A11
Targeting Non-9G4 BCR
Viability (%)
E:T Ratio
Control T cells

**Fig. S1. Design and characterization of different 9G4 CAR-T cells. (A)** Diagrams highlighting different 9G4 CAR-T cell designs. 9G4 CAR "EAAAK-CD8α Hinge" T cells comprise an EAAAK linker, CD8α hinge, CD28 transmembrane domain [TM], in addition to a CD28 costimulatory [co-stim] domain and CD3ζ signaling domain, present in all designs; 9G4 CAR "G4S-IgG4 Hinge" T cells comprise a G4S linker, IgG4 hinge, and CD28 TM; 9G4 CAR "EAAAK-IgG4 Hinge" T cells utilize an EAAAK linker, IgG4 hinge, and CD28 TM; 9G4 CAR "$(EAAAK)_3$" T cells utilize an EAAAKx3 linker, no designated hinge, and a CD28 TM; 9G4 CAR "EAAAK-CD3ζ Hinge/TM" T cells utilize an EAAAK linker, a CD3ζ hinge, a CD3ζ TM, and a CD28 co-stim domain. **(B)** Flow cytometric quantification of editing rates of engineered CAR T cells, shown in A, and control T cells (unedited, TCR KO) at four days after nucleofection. **(C)** Flow cytometric analysis showing purity of the same T cells following positive selection. Edited cells were visualized by staining for truncated NGFR (knocked into *TRAC* with CAR, expected to be positive) and CD3 epsilon (CD3, expected to be negative after *TRAC* KO). **(D)** Histograms (from left to right) showing flow cytometric staining for truncated NGFR, CD3 epsilon, and three SLE patient-derived 9G4id monoclonal antibodies (mAbs; clones 627A11, 75G12, 88F7) to show CAR expression. Median Fluorescence Intensity (MFI) for each staining is shown next to each histogram. **(E)** Summary statistics comparing MFIs of truncated NGFR staining for each 9G4 CAR-T cell and control T cells. Data are shown as mean ± SD of 4 replicates. **(F)** MFIs of monoclonal 9G4id 88F7 binding to the same set of T cells shown in E. Data are shown as mean ± SD of 2 replicates. *****P*<0.0001, ****P*<0.001; one-way ANOVA with Tukey's multiple comparison test. **(G)** Engineered 9G4 CAR-T cells were incubated with $1x10^4$ 9G4id $BCR_{627A11}$ Ramos B cells (left graph) or non-9G4 Ramos B cells (right graph) for 65 hours at different E:T ratios. The absolute number of single, live GFP+, StrepTactin XT+ B cells, as determined by flow cytometry, was used to calculate % viability for each condition. All data were normalized to the control T cell condition (E:T=0). Details of statistical analyses are provided in Dataset.

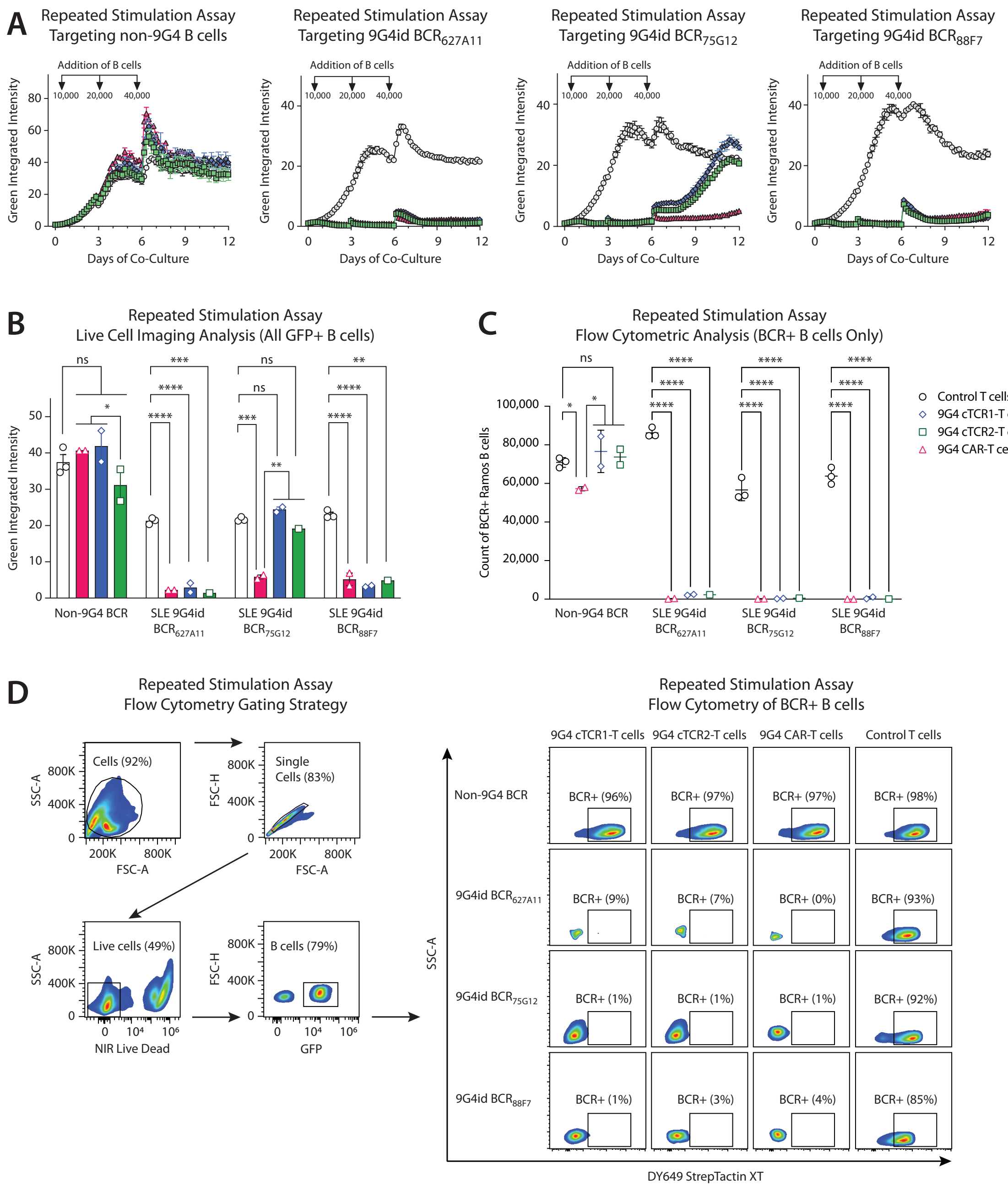


**Fig. S2. Long-term functional assessment of 9G4 synthetic immune receptor T cells in vitro. (A)** Changes in Green Integrated Intensity in RSA. 50,000 engineered T cells with 10,000 GFP-expressing Ramos non-9G4 B cells, $BCR_{627A11}$, $BCR_{75G12}$, $BCR_{88F7}$, as quantified by live-cell imaging. New Ramos B cells (but not new T cells) were added to the cocultures every 72 hours and Green Integrated Intensity was monitored for 12 days. **(B)** Quantification of green integrated intensity of total GFP+ Ramos B cells at the end of the repeated stimulation assays shown in A. Data were normalized to day 0 for each condition. ****$P$<0.0001, ***$P$<0.001, **$P$<0.01, *$P$<0.05, ns, not significant; two-way ANOVA with Tukey's multiple comparison test. **(C)** Flow cytometry quantifying B cells that express

9G4id+ BCRs (GFP+, DY649 StrepTactin XT+) at the end of the repeated stimulation assays in A-B. Data are shown as mean ± SD of technical replicates, except for panel A: Ramos $BCR_{627A11}$ x cTCR2-T cells. Comparisons are representative of n=2 independent experiments. ****$P$<0.0001, *$P$<0.05, ns, not significant; two-way ANOVA with Tukey's multiple comparison test. **(D)** Gating strategy and representative dot plots of autoreactive BCR+ and BCR- Ramos B cells at the end of repeated stimulation assays. Details of statistical analyses are provided in Dataset.

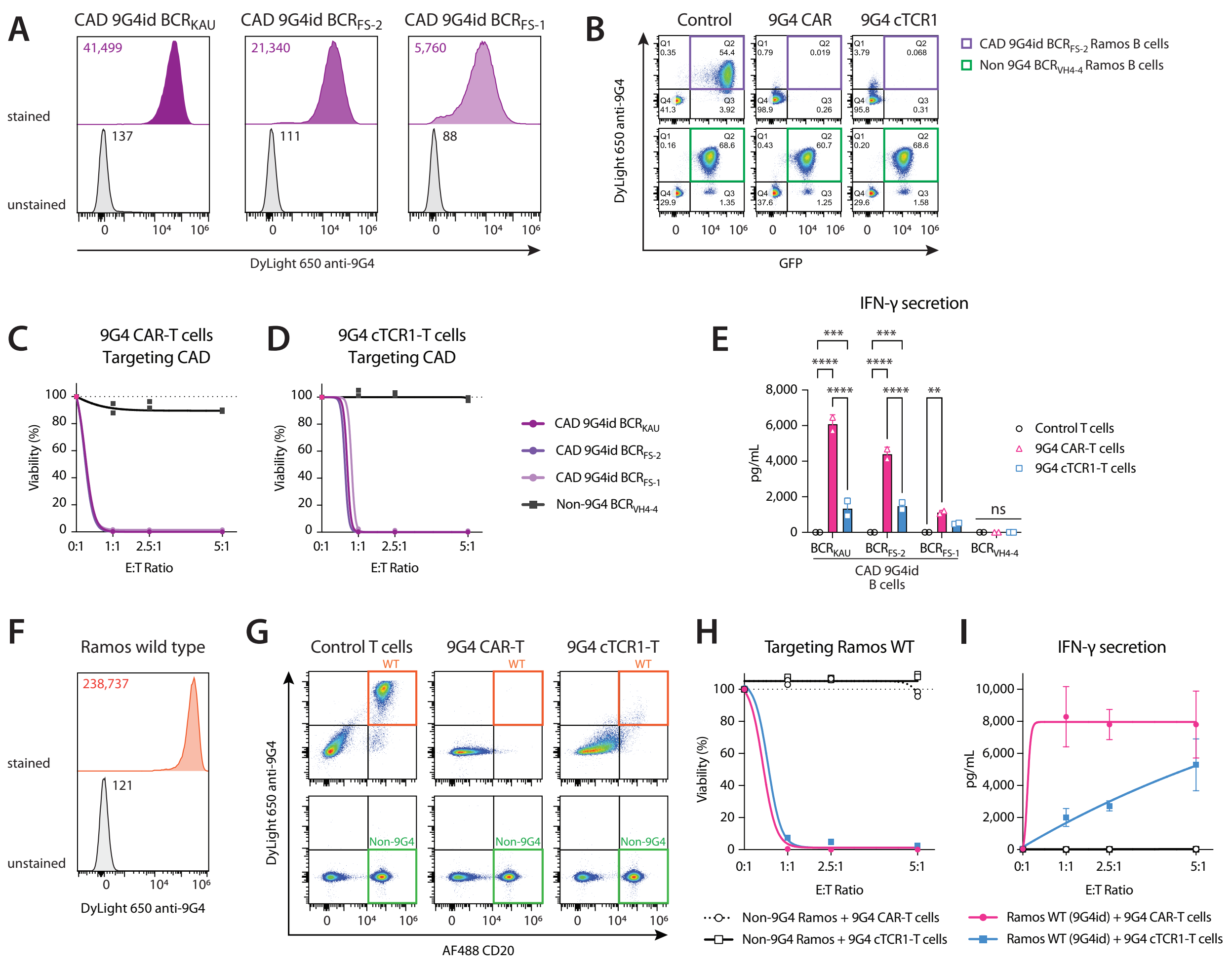


**Fig. S3. Efficacy of 9G4 CAR-T cells and 9G4 cTCR-T cells against autoreactive 9G4id B cells in cold agglutinin disease (CAD) and B cell lymphoma. (A)** Histograms showing flow cytometric staining of three 9G4id Ramos B cell lines expressing CAD patient-derived cold agglutinin BCRs (CAD 9G4id Ramos $BCR_{KAU}$, $BCR_{FS-2}$, $BCR_{FS-1}$) with an anti-9G4 antibody. Median Fluorescence Intensity (MFI) for each stained and unstained B cell clone is shown. **(B)** Representative flow cytometry plots showing the selective killing of Ramos B cells expressing the autoreactive CAD 9G4id $BCR_{FS-2}$ (top row) at the end of co-culture with either control T cells ("Control", left panel), 9G4 CAR-T cells (middle panel), or 9G4 cTCR1-T cells (right panel). 9G4id $BCR_{FS-2}$ B cells (Q2, top row) were eliminated by 9G4 cTCR1-T cells and 9G4 CAR-T cells. Ramos B cells expressing a non-9G4 BCR/antibody (VH4-4), used as an irrelevant B cell control, were not depleted under the same conditions (Q2, bottom row). Single, live cells after 3 days of co-incubation are shown (E:T ratio=5:1). **(C-D)** Percent viability of autoreactive Ramos B cells (CAD 9G4id Ramos $BCR_{KAU}$, $BCR_{FS-2}$, $BCR_{FS-1}$) or control Ramos B cells (non-9G4 BCR) at the end of co-culture with either 9G4 CAR-T cells **(C)** or 9G4 cTCR1-T cells **(D)** at different E:T ratios (E:T=5:1, $5x10^4$ edited T cells; E:T=2.5:1,

2.5x$10^4$ edited T cells; E:T=1.25:1, 1.25x$10^4$ edited T cells; 1x$10^4$ B cells). The absolute numbers of single, live GFP+, StrepTactin XT+ B cells, as determined by flow cytometry, were used to calculate % viability. All data are normalized to the control T cell condition (E:T=0). Two-way ANOVA with Dunnett's multiple comparison test. **(E)** Quantification of IFN-γ in conditioned media of co-cultures shown in (C-D), as measured by ELISA. Data are shown as mean ± SD of technical replicates. Two-way ANOVA with Tukey's multiple comparison test. **(F)** Histograms of flow cytometric staining verifying the expression of a native 9G4id BCR+ by the Burkitt lymphoma cell line Ramos RA1 (wild type, WT). **(G)** Representative flow cytometry plots showing the selective killing of WT Ramos B cells expressing 9G4id BCRs (top row) at the end of co-culture with control T cells, 9G4 CAR-T cells, or 9G4 cTCR1-T cells (top panels, from left to right). Ramos B cells expressing a non-9G4 BCR/antibody (VH4-4) were not depleted under the same conditions (bottom row). In this example, flow cytometric analysis of single, live cells after 3 days of co-incubation at an E:T ratio of 5:1 (5x$10^4$ edited T cells, 1x$10^4$GFP+ Ramos B cells) is shown. **(H)** Percent viability of Ramos B cells (WT 9G4id BCR or non-9G4 BCR) at the end of co-culture with either 9G4 CAR-T cells or 9G4 cTCR1-T cells at different E:T ratios. The absolute number of single, live GFP+, DyLight 650 anti-9G4+ or GFP+, DyLight 650 anti-9G4- B cells were used to calculate % viability. All data were normalized to the control T cell condition (E:T=0). Data are shown as mean ± SD of technical replicates; two-way ANOVA with Tukey's multiple comparison test. **(I)** Quantification of IFN-γ in conditioned media of co-cultures shown in (G), as measured by ELISA. Comparisons are representative of n=2 independent experiments. ****$P$<0.0001, **$P$<0.01, ns, not significant; two-way ANOVA with Tukey's multiple comparison test. Details of statistical analyses are provided in Dataset.

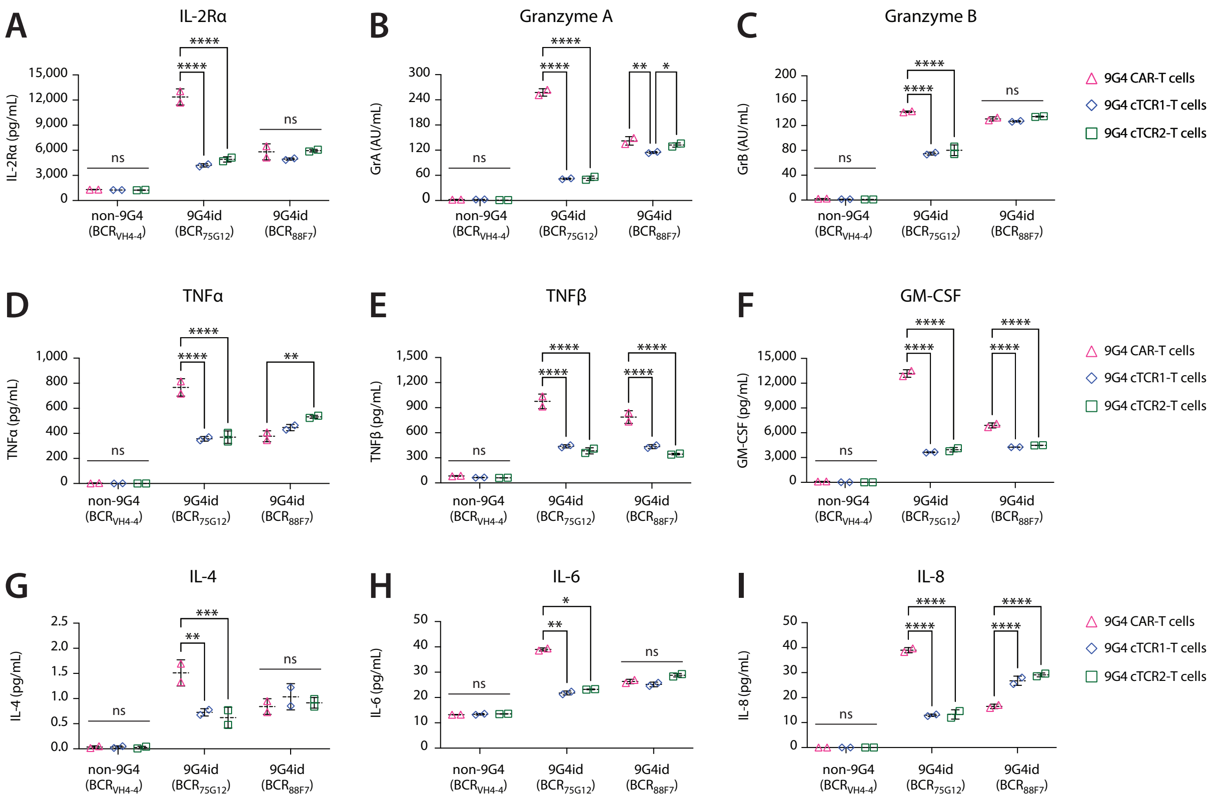


**Fig. S4. Cytokines and cytotoxic granule proteins release by 9G4 CAR-T cells and 9G4 cTCR-T cells targeting Ramos B cells. (A-I)** Quantification of cytokines (IL2Rα, TNF-α, TNF-β, GM-CSF, IL-4, IL-6, IL-8) and secreted cytotoxic granule proteins (granzyme A [GrA], granzyme B [GrB]) in co-culture supernatants of the same experiment shown in Figure 4A by U-PLEX (Meso Scale). Data are expressed as mean ± SD. ****$P$<0.0001, **$P$<0.01, *$P$<0.05, ns, not significant; two-way ANOVA with Tukey's multiple comparisons were performed. Details of statistical analyses are provided in Dataset.

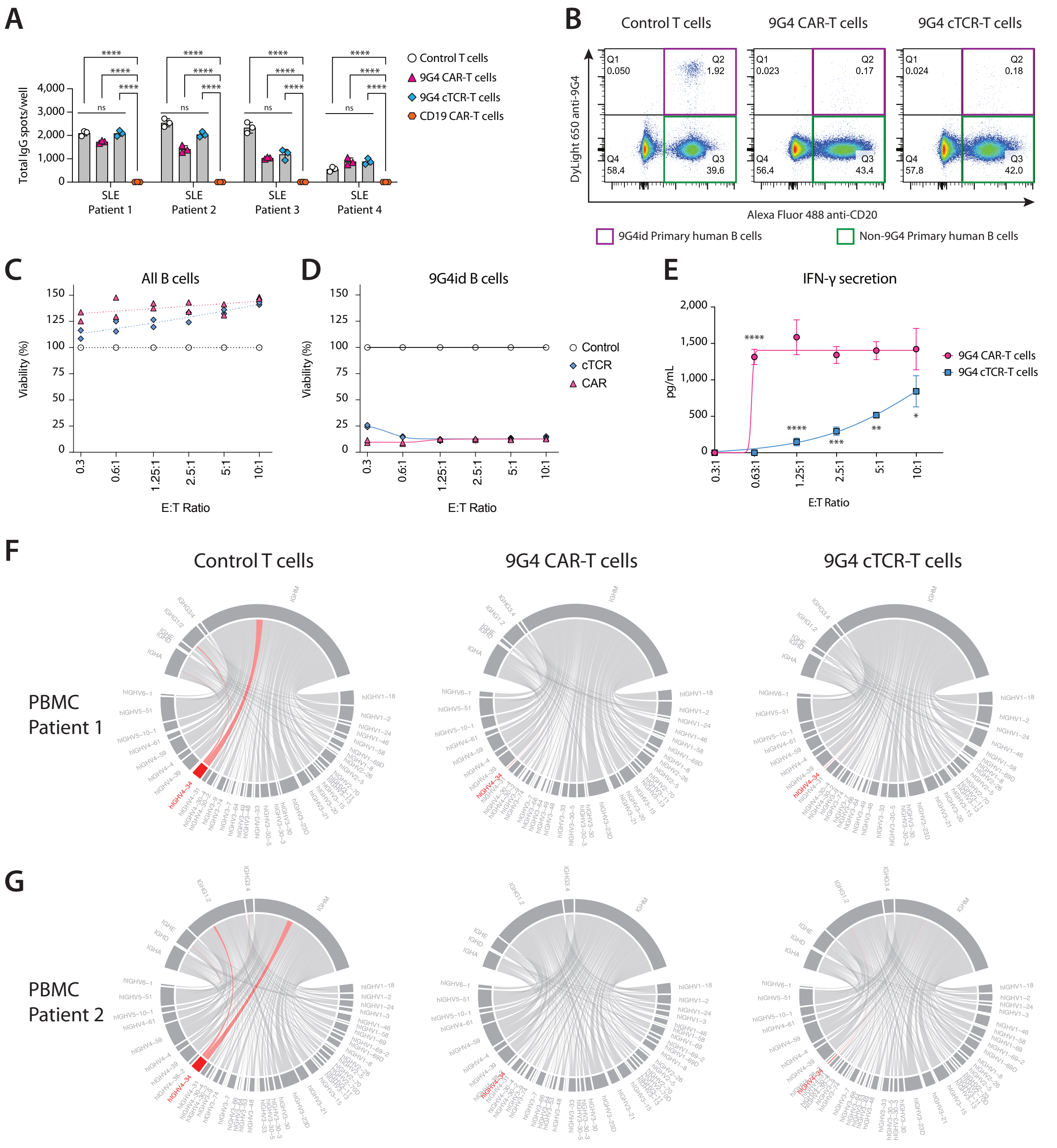


**Fig. S5. 9G4 cTCR-T cells and 9G4 CAR-T cells eliminate 9G4id B cells from patients with SLE. (A)** Quantification of B cell FluoroSpot results shown in Figure 5A showing the total number of IgG antibody secreting cell (ASC) spots per well. **(B-D)** Co-culture of isolated primary human B cells ($1x10^5$, containing ~5% 9G4id B cells) with autologous control T cells (TCR KO), 9G4 CAR-T cells, or 9G4 cTCR1-T cells for 65 hours at different effector-to-target (E:T) ratios. **(B)** Representative flow plots showing primary 9G4id B cells (CD20+, anti-9G4+; Q2) and non-9G4 B cells (CD20+, anti-9G4-; Q3) at the end of co-incubation. **(C)** Percent viability of total primary human B cells after treatment with autologous control T cells (TCR KO), 9G4 CAR-T cells, or 9G4 cTCR1-T cells. **(D)** Percent viability of primary human 9G4id B cells after treatment with autologous control T cells (TCR KO), 9G4 CAR-T cells, or 9G4 cTCR1-T cells. Data were normalized to control T cell conditions. ****$P$<0.0001, ***$P$<0.001, **$P$<0.01, *$P$<0.05; two-way ANOVA with Tukey's multiple comparison test. **(E)** Quantification of IFN-γ in co-culture supernatants of primary human B cells with autologous 9G4 CAR-T cells or 9G4 cTCR1-T cells shown in (B-D), as measured by ELISA. ****$P$<0.0001, ***$P$<0.001, **$P$<0.01, *$P$<0.05; two-way ANOVA with Šídák's correction. **(F-G)** IGHV mRNA sequencing of SLE patient PBMCs after incubation with autologous control T cells (left), 9G4 CAR-T cells (middle), or 9G4 cTCR1-T cells (right). Chord diagrams show the usage of IGHV and IGHC genes. IGHV4-34+ B cell clonotypes, which predominantly resided in the IgG1/2 and IgM compartments, were depleted after treatment with 9G4 CAR-T cells or 9G4 cTCR-T cells. Data show is for SLE patient 1 (F) and SLE patient 2 (G), respectively. Data are shown as mean ± SD of technical replicates. Comparisons are representative of n=2 independent experiments. Details of statistical analyses are provided in Dataset.

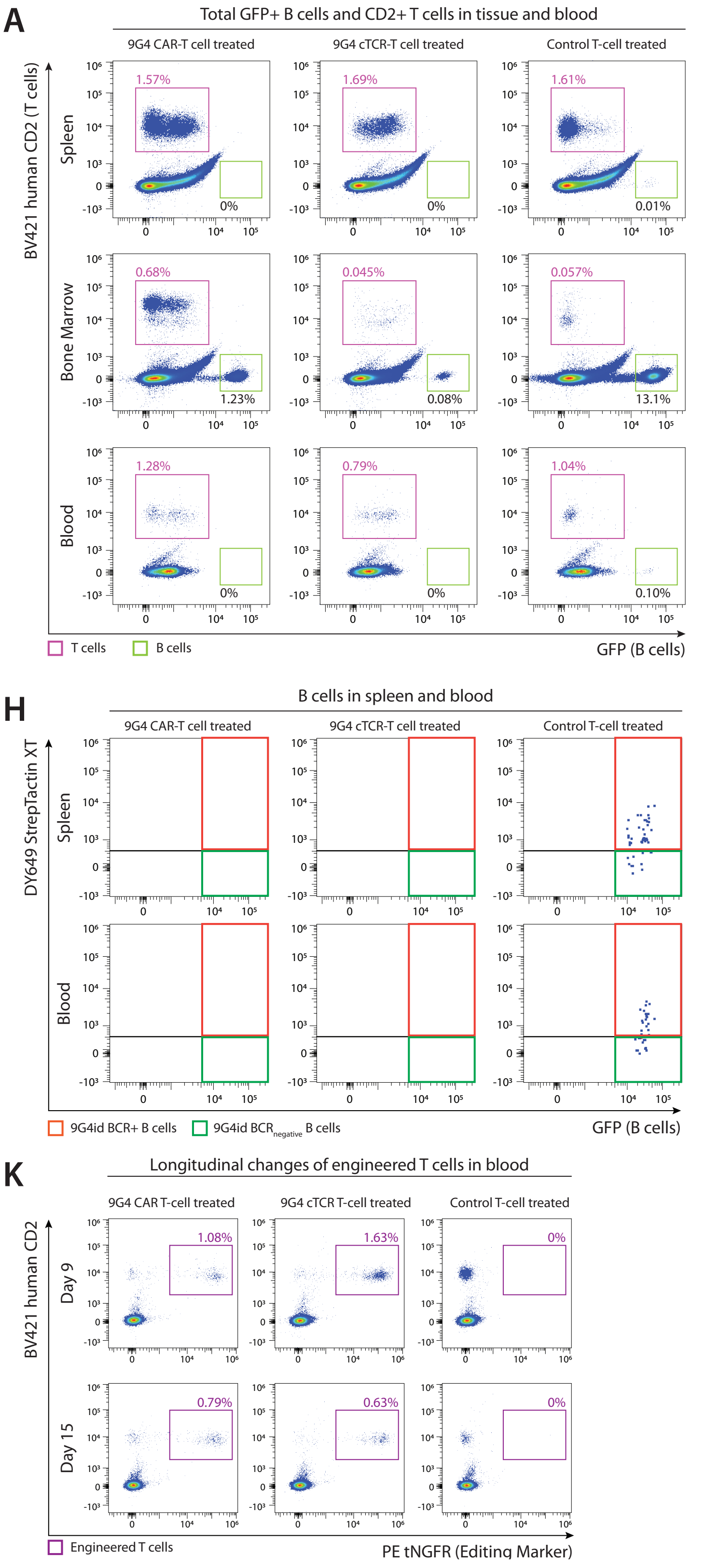

A
Total GFP+ B cells and CD2+ T cells in tissue and blood
9G4 CAR-T cell treated
9G4 cTCR-T cell treated
Control T-cell treated
BV421 human CD2 (T cells)
Spleen
Bone Marrow
Blood
GFP (B cells)
T cells
B cells
B
C
D
E
F
G
%GFP+ B cells of total splenocytes
%GFP+ B cells of total bone marrow cells
%GFP+ B cells of total nucleated blood cells
%CD2+ T cells of total splenocytes
%CD2+ T cells of total bone marrow cells
%CD2+ T cells of total nucleated blood cells
9G4 CAR
9G4 cTCR
Control
H
B cells in spleen and blood
DY649 StrepTactin XT
9G4id BCR+ B cells
9G4id BCR negative B cells
I
9G4id BCR+ B cells in Spleen
J
9G4id BCR+ B cells in Blood
BCR+ B-cell count per 10⁵ splenocytes
BCR+ B-cell count per 10⁵ nucleated blood cells
K
Longitudinal changes of engineered T cells in blood
9G4 CAR T-cell treated
9G4 cTCR T-cell treated
Control T-cell treated
BV421 human CD2
Day 9
Day 15
Engineered T cells
PE tNGFR (Editing Marker)

**Fig. S6. Flow cytometric quantification of B cells and T cells in vivo. (A)** Representative flow cytometry plots of human CD2 (T cells) versus GFP (Ramos B cells) in the spleen, bone marrow, and blood at endpoint for mice treated with 9G4 CAR-T cells, 9G4 cTCR2-T cells, or TCR KO (control) T cells. **(B-D)** Quantification of total GFP+ B cells as a percentage of total splenocytes (B), of total bone marrow cells (C), or of total nucleated blood cells (D). **(E-G)** Quantification of total human CD2+ T cells as a percentage of total splenocytes (E), total bone marrow cells (F), or total nucleated blood cells (G). **(H)** Representative flow plots of SLE 9G4id BCR+ B cells in the spleen and peripheral blood at endpoint; gates indicate 9G4id BCR+ B cells and $BCR_{negative}$ B cell subsets. **(I–J)** Quantification of 9G4id BCR+ B cell counts per $10^5$ splenocytes (I) and per $10^5$ nucleated blood cells (J). **(K)** Representative flow plots of engineered T cells (CD2+tNGFR+) at days 9 and 15 in peripheral blood. Each symbol represents an individual mouse; Data are shown as mean ± SD. One-way ANOVA with Tukey's multiple comparisons were utilized in B-G, I-J. Details of statistical analyses are provided in Dataset.

**Table S1. Patient characteristics.**

| ID | Diagnosis | Age | Gender | Treatment at visit | SLEDAI-2k at visit | Features of disease at visit |
|---|---|---|---|---|---|---|
| **Patient 1** | SLE, APS | 52 | Male | hydroxychloroquine | 2 | anti-dsDNA, anticardiolipin, anti-B2GPI antibodies, lupus anticoagulant |
| **Patient 2** | SLE, APS | 33 | Female | hydroxychloroquine, prednisone | 4 | anemia, rash/ulceration (resolving); anti-dsDNA, anti-B2GPI antibodies |
| **Patient 3** | SLE | 29 | Female | hydroxychloroquine (on admission) | 16 | Hospitalized with lupus flare: leukopenia, thrombocytopenia, anemia, pleurisy, lupus panniculitis, arthritis, myositis, ESRD due to LN (on dialysis); anti-dsDNA, anti-Sm, anti-RNP antibodies. |
| **Patient 4** | SLE, APS | 57 | Male | hydroxychloroquine | 5 | thrombocytopenia; ANA, anti-dsDNA antibodies, anticardiolipin, anti-B2GPI antibodies, lupus anticoagulant, hypocomplementemia |

**Table S2. Frequency of B cells in SLE PBMCs and autologous engineered T cell properties.**

| | **Patient 1** | **Patient 2** | **Patient 3** | **Patient 4** |
|---|---|---|---|---|
| B-cell frequency in PBMC | 7.31% | 5.99% | 5.3% | 3.78% |
| 9G4id B-cell frequency in B cells | 6.3% | 5% | 6% | 6.11% |
| 9G4 CAR-T cells Editing rate | 11.10% | 12.20% | 10.10% | 7.74% |
| 9G4 cTCR1-Tcells Editing rate | 8.77% | 12% | 9.07% | 5.56% |
| CD19 CAR-T cells Editing rate | 27.50% | 17.60% | 17.20% | 17.70% |
| 9G4 CAR-T cells Purity | 97.10% | 90% | 91.20% | - |
| 9G4 cTCR1-T cells Purity | 95% | 96.10% | 97.20% | - |
| CD19 CAR-T cells Purity | 99.20% | 95.50% | 93% | - |
| CD8+ T-cell frequency in 9G4 CAR-T cells | 22.70% | 69.10% | 35.30% | 11.40% |
| CD8+ T-cell frequency in 9G4 cTCR1-T cells | 23% | 72.50% | 36.80% | 11.10% |
| CD8+ T-cell frequency in CD19 CAR-T cells | 23.70% | 66.10% | 32.60% | 7.89% |
| CD8+ T-cell frequency in unedited T cells (Control T cells) | 27.90% | 71.80% | 55.20% | 27.50% |
| PD1 expression in 9G4 CAR-T cells (MFI) | 323 | 232 | 265 | 313 |
| PD1 expression in 9G4 cTCR1-T cells PD-1 (MFI) | 235 | 194 | 241 | 217 |
| PD1 expression in CD19 CAR-T cells PD1 (MFI) | 384 | 293 | 334 | 362 |
| PD1 expression in Unedited T cells (Control T cells) PD-1 (MFI) | 249 | 249 | 296 | 256 |

**Table S3. Nucleotide sequences of CRISPR guide RNA.**

| Name | Description | Sequence |
|---|---|---|
| Cpf1 TRAC gRNA | Cpf1 A.s. crRNA | GAGTCTCTCAGCTGGTACAC |
| Cpf1 TRBC gRNA | Cpf1 A.s. crRNA | GCCCTATCCTGGGTCCACTC |
| Cpf1 CD3G gRNA | Cpf1 A.s. crRNA | CAGGTACTTTGGCCCAGTCA |
| Cas9 TRAC gRNA | Cas9 sgRNA | AGAGTCTCTCAGCTGGTACA |
| Cas9 TRBC gRNA | Cas9 sgRNA | GGAGAATGACGAGTGGACCC |
| Cpf1-tCTS-F | HDRT amplification forward primer for Cpf1 TRAC KIs | TGGCGGGACTAGTGGCGCACAGTAAACCTGAATCTTTGGCAGTCACGACGTTGTAAAACG |
| Cpf1-tCTS-R | HDRT amplification reverse primer for Cpf1 TRAC KIs | CACCACTTCCAGCACCGCACAGTAAACCTGAATCTTTGGAGCGGATAACAATTTCACACAGG |
| huIgH296-Cas9-gsRNA | Cas9 sgRNA | GTCTCAGGAGCGGTGTCTGT |
| Cas9-tCTS-F | HDRT amplification forward primer for Cas9 TRAC KI | TGGCGGGACTAGTGGCTCTCTCTCTCAGCTGGTACACGGCAGTCACGACGTTGTAAAACG |
| Cas9-tCTS-R | HDRT amplification reverse primer for Cas9 TRAC KI | CACCACTTCCAGCACCTCTCTCTCTCAGCTGGTACACGGAGCGGATAACAATTTCACACAGG |
| Cpf1-tCTS-F | HDRT amplification forward primer for Cpf1 TRAC KI | TGGCGGGACTAGTGGCGCACAGTAAACCTGAATCTTTGGCAGTCACGACGTTGTAAAACG |
| Cpf1-tCTS-R | HDRT amplification reverse primer for Cpf1 TRAC KI | CACCACTTCCAGCACCGCACAGTAAACCTGAATCTTTGGAGCGGATAACAATTTCACACAGG |

**Table S4. Amino acid sequences of 9G4id and control BCRs.**

| BCR Name | Clone | Chain | Amino Acid sequence |
|---|---|---|---|
| CAD KAU | KAU | VH | QVHLQQWGTGLLKPSETLSLTCAVYGGSFSDYYWSWIRQPPGKGLEWIGEINDSGITNYNPSLKSRVTISVDTSKNQFSLKLSSVTAADTALYYCAREWGSGAYYPYYWGQGNLVTVSS |
| | | VL | EIVLTQSPATLSLSPGERATLSCGASQSVSSNYLAWYQQKPGQAPRLLIYDASSRATGIPDRFSGSGSGTDFTLTISRLEPEDFAVYYCQQYGSSPLTFGGGTKVEIK |
| CAD FS-1 | FS-1 | VH | QVQLQQWGAGLLKPSETLSLTCAVYGGSFSDYYWRWIRQPPGKGLEWIAEINHSGSTNYSPSLKSRVTIAVDTSKNQFSLNMNSVTATDTALYYCARGQRGSGSDYRQGAFDIWGQGTLVTVSS |
| | | VL | DIVMTQSPLSLPVTPGEPASISCRSSQSLLHSNGFNYLHWYLQKPGQSPRLLIYLGSNRASGVPDRFSGSGSGTDFTLKISRVEADDVGIYYCMQALQSPYTFGQGTKLEIK |
| CAD FS-2 | FS-2 | VH | QVHLQQWGTGLLKPSETLSLTCAVYGGSFSDYYWSWIRQPPGKGLEWIGEINDSGITNYNPSLKSRVTISVDTSKNQFSLKLSSVTAADTALYYCAREWGSGAYYPYYWGQGNLVTVSS |
| | | VL | EIVLTQSPGTLSLSPGERATLSCRASQSVSSNYLAWYQQRPGQAPRLLIYGASSRATGIPDRLSGSGSGTDFTLTINRLEPEDFAVYFCQQYGTSPRPFGQGTRLEIK |
| Non-9G4 | P2-162 | VH | QVQLQESGPGLVNPSGTLSLTCAVSGDSISSTNWWTWVRQPPGKGLEWIGEIYESGSTNYNPSLKSRLNMSVDKSKNQFSLKLRSVTAADTAVYYCARSVAGGFSFDYWGRGTLVTVS |
| | | VL | HVILTQPASVSGSPGQSITISCTGTSSDVGAYNYVSWYQQYPGKAPKVMIYEVNKRPSGVPDRFSGSKSGNTASLTVSGLQADDEADYYCSSYAGNSNSPVFGGGTKLTVL |

**Data Files S1-S9 (separate file)**

IGHV sequencing data of SLE patient PBMCs treated with anti-9G4 or control T cells

**Dataset (separate file)**

Details and extended statistical analyses for Figures and Supplementary Figures

**Movie S1-S6.**

Live-cell imaging of anti-9G4 T cells targeting 9G4id or non-9G4 Ramos B cells
